# Disulphide and sequence-encoded conformational priors guide nanobody structure prediction

**DOI:** 10.64898/2026.02.13.705647

**Authors:** Montader Ali, Mateusz Jaskolowski, Matthew Greenig, Tung H. Nguyen, Mia Crnogaj, Eva Smorodina, Aubin Ramon, Haowen Zhao, Monica L. Fernández-Quintero, Elodie Ghedin, Victor Greiff, Pietro Sormanni

**Affiliations:** Yusuf Hamied Department of Chemistry, University of Cambridge, Cambridge, UK; Department of Pharmacology, University of Cambridge, Cambridge, UK; Department of Chemical Engineering, Imperial College London, South Kensington Campus, London, UK; Department of Immunology, University of Oslo and Oslo University Hospital, Oslo, Norway; Systems Genomics Section, Laboratory of Parasitic Diseases, National Institute of Allergy and Infectious Diseases (NIAID), National Institutes of Health (NIH), Bethesda, Maryland 20894, USA; Department of Integrative Structural and Computational Biology, The Scripps Research Institute, La Jolla, CA 92037, USA

**Author notes:** {, }. { }.

## Abstract

Nanobody binding is largely governed by the HCDR3 loop, which adopts distinct placement regimes relative to the framework: compact, framework-contacting (kinked blueprint) and solvent-exposed (extended blueprint). Many nanobodies also contain additional cysteines that form non-canonical disulphide bonds, imposing covalent constraints on binding-loop conformations. Current structure predictors are typically trained and benchmarked with smooth coordinate-based objectives, so models may appear reasonable under root-mean-square deviation (RMSD), while adopting an incorrect HCDR3 blueprint or failing to recover the native disulphide connectivity, impacting paratope geometry and functional interpretation. Here, we show that the HCDR3 blueprint is predictable from sequence alone, allowing for explicit constraints during modelling. We implement these principles into NbForge, a lightweight nanobody folding model that incorporates blueprint- and disulphide-aware inductive biases and is trained with filtered self-distillation. NbForge improves recovery of HCDR3 blueprint and non-canonical disulphide formation over previous lightweight models and achieves coordinate accuracy at par to state-of-the-art, large, resource-intensive predictors, while running at sub-second inference speed. We show that using NbForge monomer models as templates further improves the success rate of predicting nanobody-antigen complexes. Together, these results motivate blueprint- and disulphide-aware benchmarks for nanobody modelling beyond RMSD, and show that appropriate inductive biases can close the performance gap to heavyweight predictors. We make the sequence classifier (NbFrame) and NbForge available for download and via a user-friendly web server.

## 1 Introduction

Nanobodies (*V*_*HH*_s) are single-domain antibodies naturally produced in camelids [1]. These molecules are becoming increasingly recognised as a promising molecular modality for a wide range of therapeutic applications following the approval of the first nanobody drug in 2019 [2, 3]. Beyond monovalent binding, *V*_*HH*_ s are genetically encodable, modular domains that can be assembled into multivalent or multispecific formats through straightforward head-to-tail fusion (typically via flexible peptide linkers), enabling compact multi-target binders and avidity engineering [4, 5]. Their small size and high stability also make them attractive targeting ligands for engineered delivery platforms (including viral nanoparticles/virus-like particles and lipid nanoparticles for nucleic-acid delivery), and as antigen-recognition domains in engineered cell therapies such as *V*_*HH*_ -based CAR-T cells [6, 7, 8, 9]. Additionally, the compact structure of nanobodies can readily be tuned for high affinity and specificity, while often retaining favourable stability and manufacturability profiles [10, 11, 12].

Deep-learning protein structure prediction has made high-throughput modelling of antibody variable domains practical, enabling structure-informed decision making earlier in discovery or engineering pipelines [13, 14, 15, 16]. In current antibody and nanobody workflows, structural models are used to triage and cluster repertoire or library hits, prioritize candidates via paratope–epitope prediction ranking, design focused mutational panels for affinity maturation, or flag developability risks (e.g., instability, aggregation propensity, and chemical liabilities) before committing to extensive experiments [17, 18, 19, 20, 21]. Across these structure-dependent applications, the key limitation is increasingly not whether a nanobody structural model can be generated, but whether it is sufficiently accurate in the regions that matter most for function – particularly the hypervariable binding loops, whose sequence and conformational diversity remains a persistent challenge for structure prediction. [22, 23, 24].

Structurally, nanobodies are single immunoglobulin variable domains in which a rather conserved *β*-sandwich framework (FR) supports three hypervariable loops, referred to as heavy-chain complementarity determining regions (HCDR1–3), that structurally define the binding region and contain most of the antigen-contacting residues (i.e., the paratope) [25]. A defining structural tendency of *V*_*HH*_ repertoires is a HCDR3 loop that is longer on average than that of conventional antibodies. This loop is also the most variable, and it frequently contributes the biggest share of antigen contacts [26], expanding molecular reach and enabling engagement of recessed or concave epitopes, but also enlarging the conformational space that structure prediction must resolve[24]. Unlike conventional antibody *V*_*H*_ domains, nanobody *V*_*HH*_ s are devoid of a paired light (L) chain [27, 28]. Consequently, the former *V*_*H*_ –*V*_*L*_ interface remains solvent-exposed in *V*_*HH*_ s and can serve as a stabilizing framework surface onto which HCDR3 folds back in kinked conformations. Consistent with this, *V*_*HH*_ frameworks exhibit adaptations at the former interface (notably increased hydrophilicity in FR2), and the absence of *V*_*L*_ reshapes the energetic balance of loop packing versus solvent exposure [29, 30, 28]. In practice, this architectural difference reduces the size of the molecule and often introduces couplings between the framework sequence and HCDR3 loop placement, such that framework regions corresponding to the former *V*_*H*_ /*V*_*L*_ interface (including FR2) can form contacts with HCDR3 residues, and in some cases also mediate contact with the antigen [24].

This coupling between framework and HCDR3 anchoring is a key determinant that structure-prediction models must capture to effectively facilitate nanobody discovery and engineering. Experimentally solved *V*_*HH*_ structures populate two broad, qualitatively distinct HCDR3 placement blueprints that differ in loop positioning and in the extent of loop–framework contacts – extended loops with no or very limited FR engagement, and kinked loops with varying amounts of FR engagement, respectively (Figure 1, Table 7) [29]. In the extended regime, HCDR3 typically projects outward into the solvent, expanding molecular reach and enabling access to recessed epitopes. In the kinked regime, HCDR3 folds back onto the domain, packing against the framework surface and creating a paratope shaped by non-local loop–framework contacts. A kinked-to-extended swap is not a small geometric drift, but a transition between alternative packing solutions that strongly affects the paratope conformation, leading to different binding-interface geometries. Therefore, correct HCDR3 blueprint prediction has important implications for downstream tasks, including nanobody-antigen docking, paratope and epitope prediction, and mutation design.

**Figure 1.**
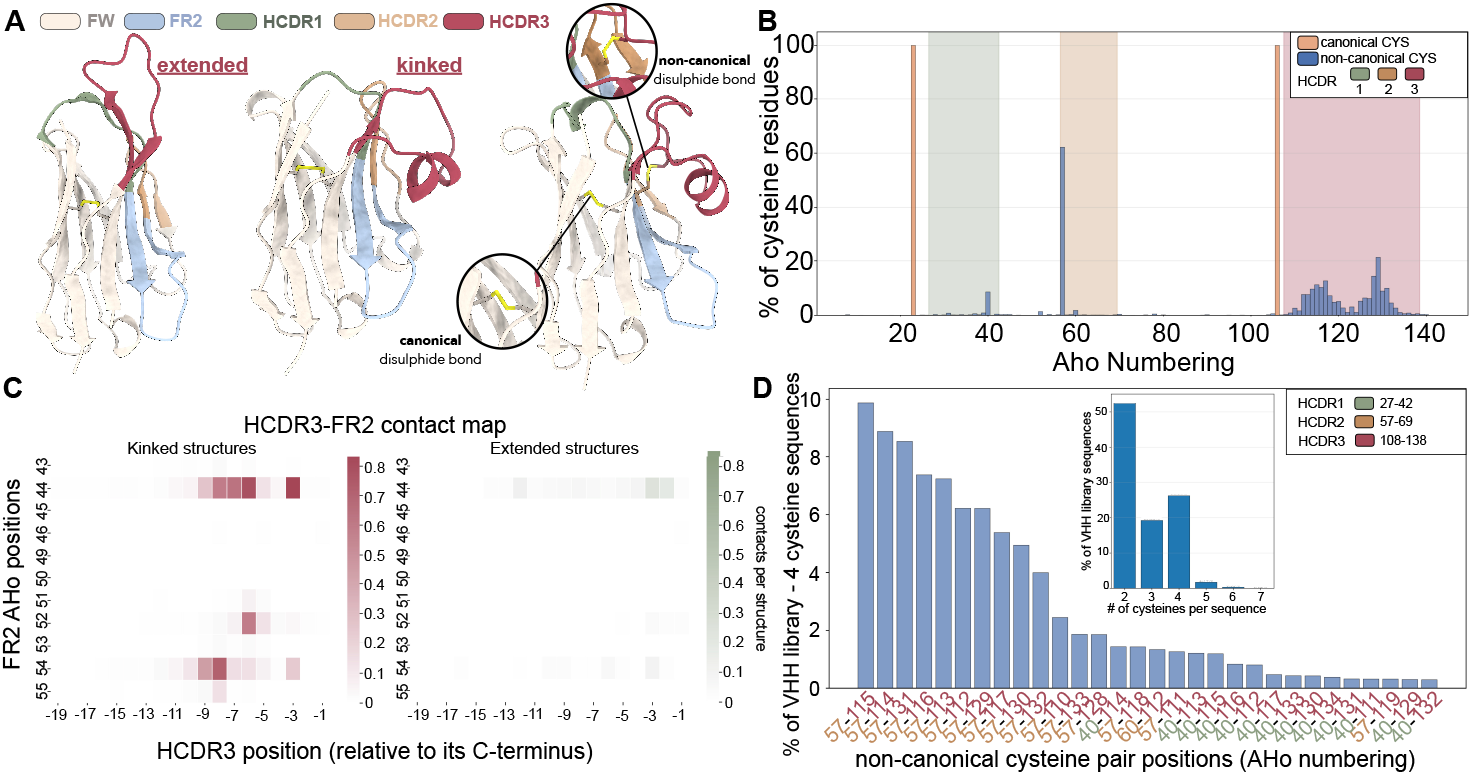
Structural and sequence features associated with kinked versus extended *V*_*HH*_ HCDR3 conformations. (**A**) Representative *V*_*HH*_ structures with an extended HCDR3 (left) and a kinked HCDR3 (centre and right). Framework (FW), framework region 2 (FR2), and HCDR1–3 are colour-coded (legend); the canonical intradomain disulphide bond is indicated and an example non-canonical disulphide involving HCDR3 is highlighted (Inset). (**B**) Distribution of cysteine residues of *V*_*HH*_ sequences in AHo numbering from a *V*_*HH*_ library of sequences [36], separating canonical and non-canonical cysteines; shaded bands denote HCDR1–3 positions. Mean HCDR3–FR2 contact maps for kinked, *n* = 481 (left), and extended, *n* = 346 (right), *V*_*HH*_ PDB structures (labelled by NbFrame), with HCDR3 indexed relative to its C terminus and FR2 positions shown in AHo numbering. (**D**) Frequencies of non-canonical cysteine-pair positions (AHo numbering) among *V*_*HH*_ library sequences [36] containing four cysteines; inset shows the distribution of total cysteine counts per sequence across the whole library.

The conformational diversity of HCDR3 reflects, in part, how *V*_*HH*_ framework features shape loop anchoring against the domain versus projection into solvent [24, 31]. While the blueprint conformations have been assigned from experimentally solved structures using geometric or contact-based criteria, structure-based definitions are not directly usable at the point where they are most needed: when reasoning about large sets of sequences that do not yet have structures (e.g., during library assessment, developability screening, or dataset construction for model training and benchmarking). In these sequence-only, discovery-scale settings, candidate triage often relies on readily computable features such as HCDR3 length distributions or cysteine motifs, which do not explicitly encode the global placement of HCDR3 relative to the framework [32, 33]. Taken together, these considerations motivate the need for a sequence-level, *V*_*HH*_ -specific prior over the HCDR3 blueprints.

A second, largely orthogonal determinant of nanobody paratope arrangement is the presence of additional, non-canonical intradomain disulphide bonds. Beyond the conserved Ig-domain disulphide, a substantial fraction of camelid *V*_*HH*_ s carry extra cysteines that form cross-links most commonly tethering HCDR3 to HCDR2, but also linking HCDR3 to HCDR1 or other framework positions near the former *V*_*H*_ –*V*_*L*_ interface (e.g., FR2), and in some cases forming intra-HCDR3 disulphides (Figure 1, Table 2) [34, 31, 12]. These bonds are frequently discussed as evolutionary and engineering solutions to the expanded conformational freedom introduced by long HCDR3 loops; analysis of the llama wild-type *V*_*HH*_ repertoire shows that over 25% of sequences contain non-canonical disulphide bonds involving CDR3 cysteines and germline-encoded cysteines at variable positions, indicating a camelid-specific mechanism of conformational preselection and stabilization that expands paratope geometry and antigen-binding diversity [35]. Indeed, by covalently constraining loop topology, they can rigidify and pre-organize binding-competent conformations, modulate loop–framework packing, and often increase structural robustness (with context-dependent effects on affinity and stability) [12, 31, 30]. Accurately capturing non-canonical disulphide bridges is also directly relevant to developability and engineering decisions. Empirically, these bonds are often coupled to thermal stability, refolding behaviour, and aggregation propensity [30, 31]. In practical discovery settings, accurate prediction of disulphide constrained conformations is important not only for structural interpretation, but also for prioritizing sequences whose folds are likely to be stable and functionally deployable.

**Table 1:** NbFrame classifier performance on held-out test structures (*n* = 100).

| Classifier | ROC-AUC | PR-AUC | Accuracy | Features |
| --- | --- | --- | --- | --- |
| Structure-based | 0.994 | 0.995 | 94.0% | 6 |
| Sequence-based | 0.939 | 0.956 | 86.0% | 20 |

**Table 2:** Summary statistics of *V*_*HH*_ datasets used in this study, including mean HCDR3 length, frequency of non-canonical disulphide bonds (NCDB), and frequency of kinked HCDR3 conformations. Note: the kinked and extended evaluations are performed using the NbFrame sequence and structure classifier for the library dataset and PDBs, respectively, and the NCDBs from the library sequences are assumed to form in all sequences with 4 and 5 cysteines.

| Dataset | Size (N) | HCDR3 length | NCDB (%) | Kinked (%) |
| --- | --- | --- | --- | --- |
| SAbDab (all $V_{HH}$ chains) | 3124 | $14.7 \pm 4.5$ | 17.0 | 69.6 |
| SAbDab (unique $V_{HH}$ sequences) | 892 | $14.4 \pm 4.3$ | 13.9 | 68.6 |
| $V_{HH}$ Library | $2.22 \times 10^7$ | $15.2 \pm 4.5$ | 26.3 | 61.8 |
| NanobodyBuilder2 test set | 30 | $14.0 \pm 4.5$ | 2.5 | 80.0 |
| NbForge test set | 47 | $17.8 \pm 4.3$ | 31.3 | 62.5 |

Taken together, the kinked/extended blueprints and non-canonical disulphide bridges highlight that nanobody loop modelling is not purely a continuous optimization problem over backbone coordinates. Instead, it is a mixed inference problem: the structure must satisfy a small number of discrete features (HCDR3 blueprint and disulphide-bridge formation), while simultaneously resolving the continuous degrees of freedom within these constraints. This observation motivates approaches that incorporate sequence-level priors over HCDR3 placement and explicit geometric objectives for disulphide-constrained topologies, as well as benchmarks that interrogate these discrete commitments directly, especially in the long-loop, cysteine-rich blueprints consequential for nanobody engineering.

We therefore introduce NbFrame, a lightweight classifier that calculates the probability of the HCDR3 adopting a kinked or extended blueprint directly from sequence, and NbForge, a lightweight, all-atom *V*_*HH*_ structure prediction framework that directly incorporates nanobody-relevant features into both training and evaluation. NbForge is trained using three complementary strategies: (i) we expand structural supervision by leveraging variable heavy chains from conventional Fv antibodies alongside curated *V*_*HH*_ structures to better cover the HCDR3 length and topology distribution; (ii) we introduce an explicit disulphide-geometry objective to encourage physically plausible non-canonical disulphide formation (including SG–SG and C*β*–C*β* distance terms, and associated angular/dihedral constraints); and (iii) we scale learning through filtered self-distillation on a large naïve *V*_*HH*_ sequence library, folding only high-confidence kinked and extended sequences and retaining only predictions that satisfy disulphide topology, avoid bond violations, and remain consistent with NbFrame-implied HCDR3 placement. The resulting synthetic set is then used to train the model and is complemented by final training/fine-tuning on experimentally solved variable-heavy chains, yielding a fast, HCDR3 blueprint and disulphide bridge constrained predictor designed for discovery-scale use.

## 2 Results

### Data curation as a prerequisite for nanobody-specific learning

A central practical constraint in nanobody structure prediction is that the available set of experimentally solved *V*_*HH*_ structures remains small and heterogeneous in quality (Table 2). We therefore began by constructing a training set from the Structural Antibody Database (SAbDab) [37], which explicitly stratifies the nanobodies by HCDR3 length, kinked and extended, as well as the presence of a non-canonical disulphide bridge. Starting from antibody-containing PDB entries, we separated conventional antibodies from nanobodies, isolated each variable heavy chain as a standalone chain-level sample (both *V*_*H*_ and *V*_*HH*_), and then removed structures containing missing residues within the heavy-chain core. Missing internal residues introduce silent label noise where the model is asked to learn a fold that is not fully specified by the coordinates, and downstream uncertainty estimate scores can be falsely attributed to loop modelling accuracy rather than incomplete supervision [38]. This procedure resulted in a consistent set of structures of antibody heavy chain variable domains (*V*_*HH*_ : *N*_*sequences*_=892, *N*_*structures*_=3128, *V*_*H*_ : *N*_*sequences*_=3672, *N*_*structures*_=13,261).

### Development of NbFrame as a structure- and sequence-based HCDR3 classifier

Given the functional importance of HCDR3 conformational blueprint, we sought to identify structural features that distinguish kinked from extended conformations in *V*_*HH*_ domains. A comprehensive feature set capturing multiple aspects of the conformational geometry surrounding the HCDR3 and the FR2 regions was used to classify the conformations of the PDB structures. We selected six features based on biological rationale and building on previous studies: backbone angles at both the N-terminal (*α*_*N*_, *τ*_*N*_) and C-terminal (*α*_*C*_, *τ*_*C*_) stems of the HCDR3, contact density between HCDR3 and FR2 (number of atomic contacts normalized by HCDR3 length), and relative solvent accessibility (RSA) of key FR2 hydrophobic positions (AHo positions 44 and 54) [29, 26].

To establish ground-truth labels, we performed Gaussian Mixture Model (GMM) clustering in the six-dimensional feature space, revealing two geometrically well-separated clusters (Figure S2; see Methods). We sampled 50 structures from each cluster, including both cluster cores and boundary cases, and classified each by manual inspection. This yielded 60 extended and 40 kinked labels. The imbalance reflects that GMM clustering captured geometric patterns primarily linked to angles, whereas the structural definition of kinked conformations additionally requires physical contacts with FR2 (see Methods).We trained a logistic regression classifier on these 100 manually labelled structures, representing each input with the six features described above. The model achieved ROC-AUC of 0.99 and accuracy of 94.0% on the held-out test set (*n* = 100) (Figure S3). Feature coefficients revealed that contact density (+1.86) and FR2 RSA (*−*1.16) were the dominant predictors (Figure S4), consistent with the functional role of kinked conformations in shielding FR2 hydrophobic residues.

As kinked and extended HCDR3 conformations are associated with characteristic framework sequence patterns, we identified framework positions where specific amino acids showed statistically significant enrichment in kinked versus extended structures (Fisher’s exact test, *p* < 0.05, minimum 30 observations). From these candidates, we selected 20 hallmark features with the strongest discriminative signal at biologically relevant positions (Figure S5): FR1 (positions 12, 15, 17), FR2 (positions 44, 51, 54, 56), and FR3 (positions 85, 103, 107). Position 107, adjacent to the conserved Cys106 at the HCDR3 junction, contributed multiple hallmarks, while positions 44 and 54 correspond to the classic FR2 hydrophobic residues that interact with kinked HCDR3 loops (Figure 1). Each sequence was encoded by the log_2_ fold-change enrichment value of each of these 20 hallmarks, when the sequence contained that specific amino acid at that position (Table 7).

We trained a logistic regression model from the sequences of 829 labelled structures, deliberately restricting features to framework positions to avoid potential bias from hypervariable CDR sequences or unusual CDR motifs found in engineered or synthetic *V*_*HH*_ s. Evaluated on the same 100 held-out structures with manually validated labels, the sequence classifier achieved ROC-AUC of 0.94 and accuracy of 86.0% (Table 1, Figure S3 and S9).

### Building a self-distillation dataset of nanobody structures from filtered immune repertoires

*V*_*HH*_ repertoires span an enormous sequence space (often quoted as ∼ 10^12^–10^15^ possible antibodies), and the long HCDR3 loops that are common in single-domain antibodies further amplify conformational diversity and consequent modelling uncertainty [22, 39, 40, 41]. Experimentally determined nanobody structures sample only a small fraction of this diversity (892 unique sequences in our curated set), limiting the sequence–structure coverage that can be learned directly from ground-truth data. To expand effective coverage while retaining modelling fidelity, we therefore adopted a self-distillation strategy: starting from a large naïve *V*_*HH*_ sequence collection (22 million sequences) [36], we applied two sequential filters to enrich for sequence diversity while favouring sequence features and structural filters consistent with accurate structure prediction (Figure 2D).

**Figure 2.**
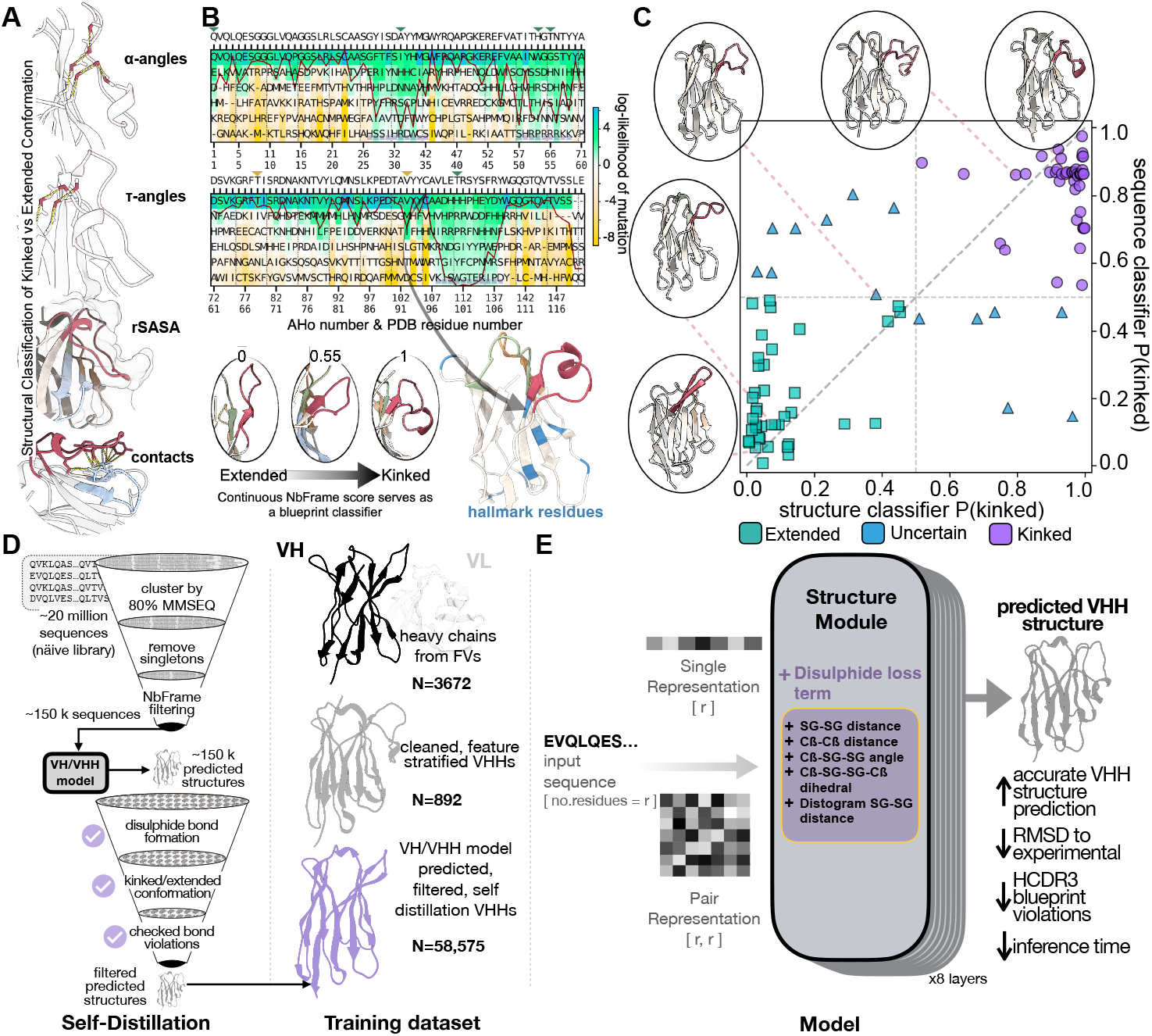
NbFrame structure and sequence classifier, creation of the self-distillation dataset, and architecture of the NbForge model. **(A)** Structural definition of the kinked/extended HCDR3 blueprints using geometric descriptors; these features provide a continuous NbFrame score that classifies blueprint states. **(B)** Sequence-based classification using framework hallmark residues via log fold-change values of kinked vs extended blueprints (see Table 7) to provide a kinked probability score. **(C)** Agreement between sequence-based and structure-based NbFrame classifier scores-with extended, kinked, and ambiguous cases shown with representative structures. **(C)** Overview of the NbForge *V*_*HH*_ self-distillation sequence and structure filtering steps. Sequences from the *V*_*HH*_ library are first clustered by MMSEQ2 at 80% similarity (the first sequence from each cluster is taken); sequences that were not clustered (singletons) were filtered out. NbFrame’s sequence classifier was used to remove sequences classified as uncertain. A *V*_*H*_ /*V*_*HH*_ trained model (see Methods) folded the filtered sequences. These predicted structures were then filtered for structural violations, incongruent HCDR3 blueprint, or missed expected NCDB, and the remaining structures were used to train the final model together with the experimental *V*_*H*_ and *V*_*HH*_ structures. **(E)** NbForge architecture for unbound *V*_*HH*_ structure prediction. During the forward pass (training and inference), the input sequence is embedded into per-residue (‘single’/node) features, and residue-residue (‘pair’/edge) features are initialised with relative positional encodings (L×L). Single and pair features are then iteratively refined within an invariant point attention (IPA)-based geometry module (AF2 structure-module-style update stack; same IPA-based architecture family used in NanoBodyBuilder2), which updates residue rigid frames to produce 3D coordinates and per-residue confidence (pLDDT head not shown). An additional ‘disulphide objective’ loss term is used during training to favour physically valid disulphide bridges.”

First, we clustered sequences using MMseqs [42] at 80% full-sequence identity and removed singletons, yielding approximately 650,000 non-redundant sequences. Second, to focus specifically on sequences with clear HCDR3 conformational blueprint markers, we retained only those predicted with high confidence to be kinked (NbFrame score 0.99) or extended (NbFrame score 0.20), intentionally removing ambiguous intermediates. Although this filtering step narrows the scope of the synthetic set, it provides a principled way to discard low-confidence models that violate the expected structural blueprint, yielding a higher-quality set for downstream training. After this stage, we obtained 154,115 sequences, which were folded at scale with a *V*_*H*_ /*V*_*HH*_ -trained version of the model (Table 2; see Methods). Briefly, this *V*_*H*_ /*V*_*HH*_ teacher model is a lightweight, structure-module–only predictor derived from ABodyBuilder3/AlphaFold2; it retains the AlphaFold2 structure module and pLDDT head, rather than the full Evoformer/MSA/template pipeline (see Methods; Figure 2E). We augment the base coordinate objectives with an explicit disulphide-bridge loss that supervises cysteine pairing and geometry.

To reduce the occurrence of inaccuracies, which can later bias the training when such modelled structures are used for self-distillation, we applied a stringent, nanobody-specific filtering strategy. First, we removed predictions with mispaired disulphide bonds (see Methods). Second, we filtered out structures whose predicted HCDR3 blueprint was inconsistent with the NbFrame-evaluated blueprint based on the structure classifier (Table 1, Figure 2, see (Methods). Furthermore, we applied TopModel (amide planar bond) [43] and peptide bond filtering that removed structures with backbone violations. Finally, we required a minimum mean HCDR3 pLDDT (0.7) as a coarse confidence criterion, ensuring that retained structures were not only topologically plausible but also internally self-consistent under the model’s own uncertainty estimates.

In summary, this curation strategy yields a synthetic self-distillation set comprising 58,575 structural models, enriched for sequences with clear kinked/extended commitments and for predictions that satisfy disulphide-bridge formed structures with loops whose conformation is predicted with high pLDDT confidence. These models are generated using a *V*_*H*_ /*V*_*HH*_ -trained NbForge teacher check-point (the same lightweight AlphaFold2 structure-module + pLDDT architecture, augmented with our disulphide-geometry objective-see Methods) trained on the curated set of experimentally solved *V*_*H*_ and *V*_*HH*_ domains. We use this teacher to fold the 154,115 blueprint-confident library sequences at scale, then retain only predictions that pass the NCDB-specific topology, blueprint-consistency, and bond-violation filters described above and in [44]. The resulting 58,575-model set provides synthetic supervision for self-distillation: we retrain the NbForge architecture under blueprint and disulphide constraints on these filtered models, and then fine-tune on experimentally solved heavy chains to obtain the final predictor.

### Dataset composition and benchmark design

To contextualise the conformations represented in our training, synthetic, and evaluation data, we summarise key HCDR3 and cysteine-linked properties across datasets in (Table 2). Experimental *V*_*HH*_ structures in SAbDab exhibit a mean HCDR3 length of approximately 14 residues (14.7±4.5 for all chains, 14.4±4.3 after deduplication), with non-canonical disulphide bonds present in a minority of cases (17.0% and 13.9%, respectively) and a strong skew toward kinked conformations (69.6% and 68.6%). In contrast, the naïve 22 million *V*_*HH*_ sequence library we use to approximate repertoire-scale diversity is shifted toward both longer loops (15.2±4.5) and a higher prevalence of non-canonical cysteine patterns (26.3%), with a modest reduction in kinked frequency (61.8% labelled via NbFrame).

We therefore designed our held-out evaluation test set of *V*_*HH*_ structures to reflect these statistics. The NanobodyBuilder2 (NBB2) test set closely mirrors SAbDab in mean HCDR3 length (14.0±4.5) but contains very few non-canonical disulphide bonds (2.5%) and is strongly enriched for kinked HCDR3s (80.0%), limiting its ability to stress test disulphide-constrained loop topologies and a balanced blueprint conformation. By contrast, our NbForge test set is deliberately shifted toward longer HCDR3 loops (17.8±4.3) and substantially higher non-canonical disulphide prevalence (31.3%), while maintaining a more balanced kinked/extended composition (62.5% kinked) and with most sequences displaying up to 50% HCDR3 residue differences to the training set (Figure S1). Moreover, we ensured that all of our test-set structures were released after December 2022 (past the AF3/Boltz1 cut-off date of September 2021 and that of NBB2 of July 2021), which - together with the high diversity from the training set (i.e., from all other *V*_*HH*_ structures) - also makes it a good benchmark set for these modelling tools. To ensure no data-leakage, we removed PDBs with homologous sequences to this test set, and show that our test set’s HCDR3 sequences are at most ∼ 60% similar to the training/validation set (Figure S1). This test-set design enables evaluation not only of coordinate accuracy, but also of whether models recover the structural features that determine the *V*_*HH*_ loop conformation - which are under-represented in standard benchmarks (HCDR3 blueprints and non-canonical disulphide bridges).

### Coordinate accuracy on the NbForge test set

NbForge is built as a lightweight model using only the structure module and pLDDT module from AlphaFold2 (see Methods, Figure 2D). The structure-module focused architecture for a nanobody folding model was initially used by the NanoBodyBuilder2 (NBB2) model, which displayed competitive RMSDs against AlphaFold2 nanobody structure prediction, at a fraction of the inference time [45], and is widely used in *V*_*HH*_ discovery campaigns [46]. Larger diffusion-based models have since shown to be the more accurate models in predicting *V*_*HH*_ s backbones [23, 47, 24]; these large models rely on sophisticated architectures, that demand extensive GPU resources, MSA inputs, with some models also accepting template structures, and are substantially more costly in terms of inference time (Table 3). We therefore evaluate our NbForge model against the lightweight NBB2 model, as well as the diffusion-based AF3 (best-in-class) and Boltz1 heavy models (open-source). Boltz2 and other recently released models could not be compared due to large overlap between their training sets and our test set, but Boltz2 has already been shown to perform similarly to Boltz1 on *V*_*HH*_ s [48].

**Table 3:**
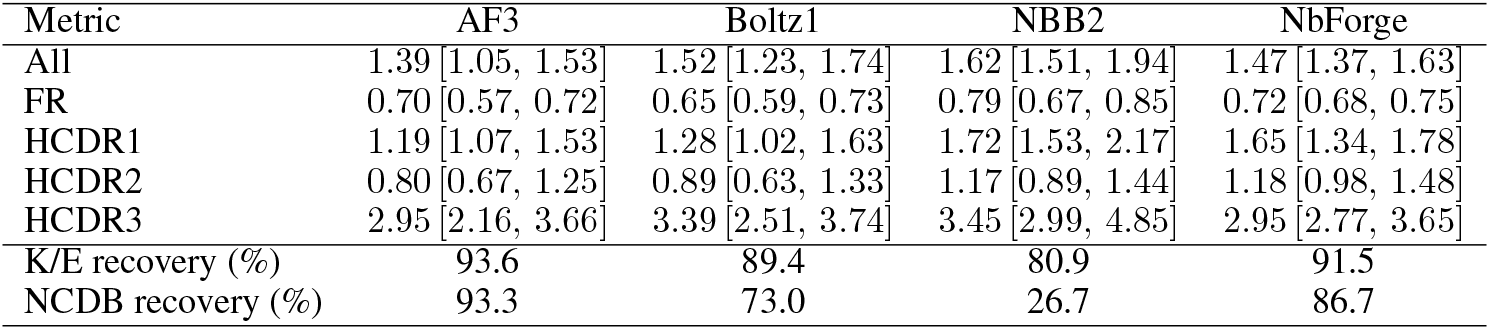
Coordinate accuracy and feature recovery on the NbForge test set. Backbone (CA, N, CO, O) RMSDs are reported as median with bootstrap 95% confidence intervals (CI) in Å (*n* = 47) after framework-region superimposition. Confidence intervals were estimated using non-parametric bootstrap resampling with 10,000 resamples. Kinked/extended (K/E) correctness is computed over all test structures, while non-canonical disulphide bond (NCDB) recovery is computed over the subset containing a non-canonical disulphide bond (*n* = 15), see Methods.

Because each model is evaluated on each *V*_*HH*_ in the same test set, we report (i) an overall repeated-measures test for model effects (Friedman) and (ii) planned paired post hoc comparisons (Wilcoxon signed-rank) focusing on NbForge versus the other models. NbForge improves significantly over NBB2 for both all-backbone RMSD and HCDR3 RMSD (*p* < 0.001 < paired Wilcoxon for both-Table S1), with large paired effect sizes (rank-biserial correlation magnitudes *∥r*_*rb*_*∥ ≈* 0.65 and |*r*_*rb*_| ≈ 0.63). Against the diffusion-based large models, NbForge is competitive on HCDR3: median HCDR3 RMSD is 2.95 Å for NbForge versus 2.95 Å for AF3 and 3.39 Å for Boltz1 (all 95% Cis overlap), and these differences are not statistically distinguishable under paired testing (NbForge vs AF3: *p* = 0.49; NbForge vs Boltz1: *p* = 0.90). However, AF3 and Boltz1 show significantly lower framework median-RMSD (0.70 Å and 0.65 Å versus 0.72 Å) (Table 3,S1), suggesting that while NbForge matches the large model’s performance on the most variable loop region, some global framework accuracy remains achievable through broader priors or MSAs, for example. Notably, in nanobodies the dominant sources of functional specificity and modelling uncertainty are concentrated in the hypervariable CDR loops, especially HCDR3, so parity in HCDR3 loop accuracy and constraint satisfaction is typically more consequential for downstream utility and design than sub-Å differences in the comparatively rigid framework.

### HCDR3 kinked/extended blueprint recovery

We next evaluated whether each method recovers the correct global HCDR3 blueprints by matching each predicted structure to its corresponding experimental chain and comparing the loop placement using NbFrame’s structural classifier definition (see Methods). All four models achieve high conformation recovery on this benchmark (Table 3), with AF3 performing best at 44/47 (93.6%), NbForge recovering 43/47 (91.6%), followed by Boltz1 at 42/47 (89.4%), and finally NBB2 at 38/47 (80.8%) (Figure 4A). The overall near-ceiling performance across models suggests that the kinked/extended distinction is strongly encoded by sequence features and can be learned reliably even from comparatively limited *V*_*HH*_ structural data, supported by the high PR-AUC of the NbFrame sequence classifier, which was built entirely from residue frequencies.

**Figure 3.**
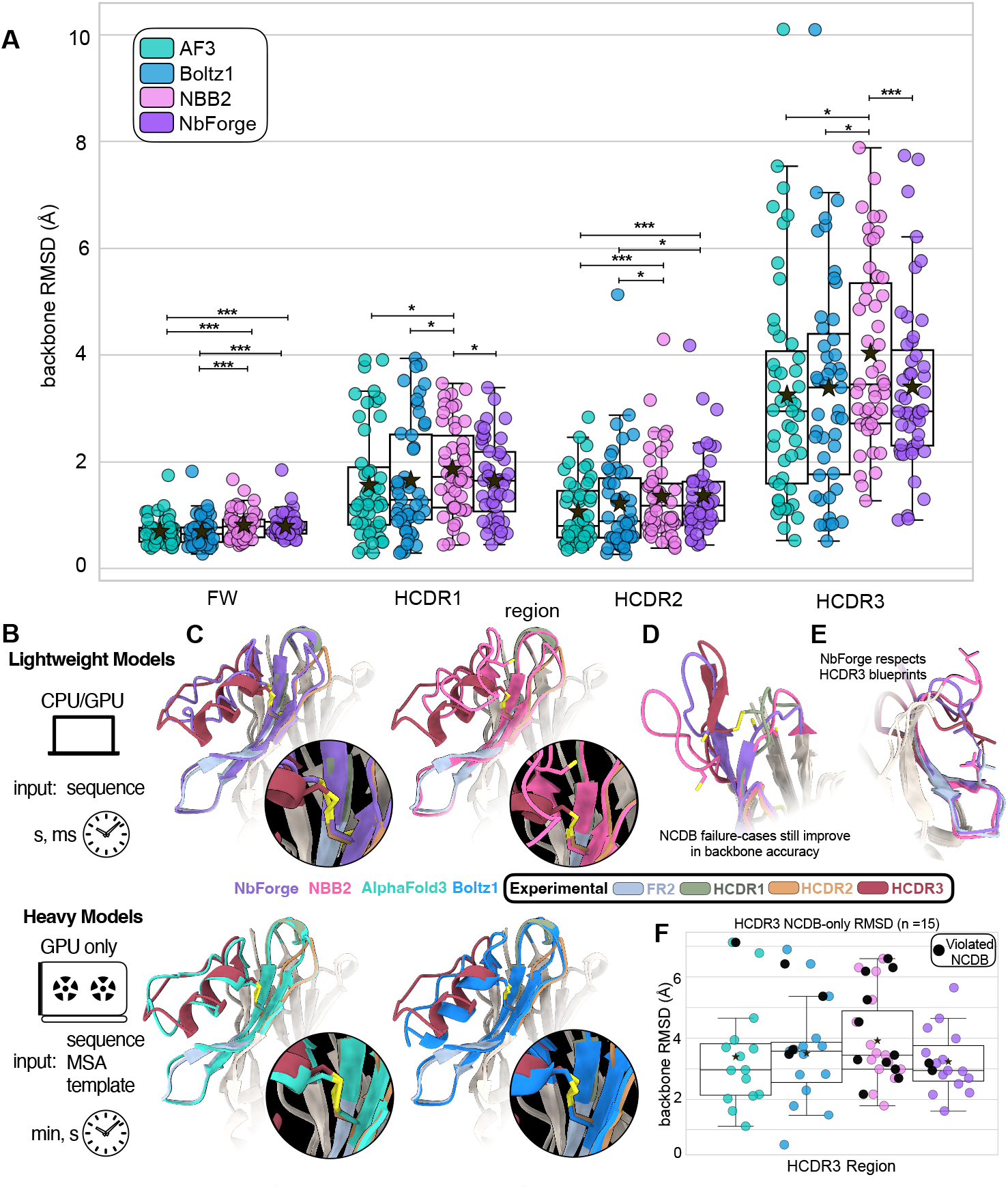
Nanobody structure prediction accuracy, HCDR3 blueprint fidelity, and NCDB formation. **(A)** Swarmplot of full-backbone RMSD to experimental structures in the held-out test set (*n* = 47) calculated following framework-region superimposition across different *V*_*HH*_ regions (x-axis) for AlphaFold3 (AF3), Boltz1, NBB2, and NbForge (legend). Boxes represent the first and third quartiles of the distribution, whiskers represent the 1.5 interquartile range, the horizontal line is the median and the star marker the average. Brackets denote Wilcoxon pairwise statistical comparison, where *, **, and *** denote *p* < 0.05, *p* < 0.01, and *p* < 0.001, respectively; see Table S1 for details. **(B)** Inference models compared in this work: “lightweight” single-sequence predictors versus “heavy” predictors. **(C)** Representative example (PDB: 9g1y) to highlight the influence of correct disulphide-bridge prediction on backbone accuracy, with close-ups highlighting loop placement and disulphide geometry. **(D)** Case study (PDB: 8z8v) highlighting NCDB-recovery failure cases that still show improved backbone accuracy among lightweight models (NbForge vs NBB2). **(E)** Case study showing that NbForge recapitulates experimental HCDR3 kinked-blueprint contacts. **(F)** Like A but only for the HCDR3-region of the NCDB-containing subset (*n* = 15), with models violating the non-canonical disulphide constraint indicated in black; values are reported in Table 4.

**Figure 4.**
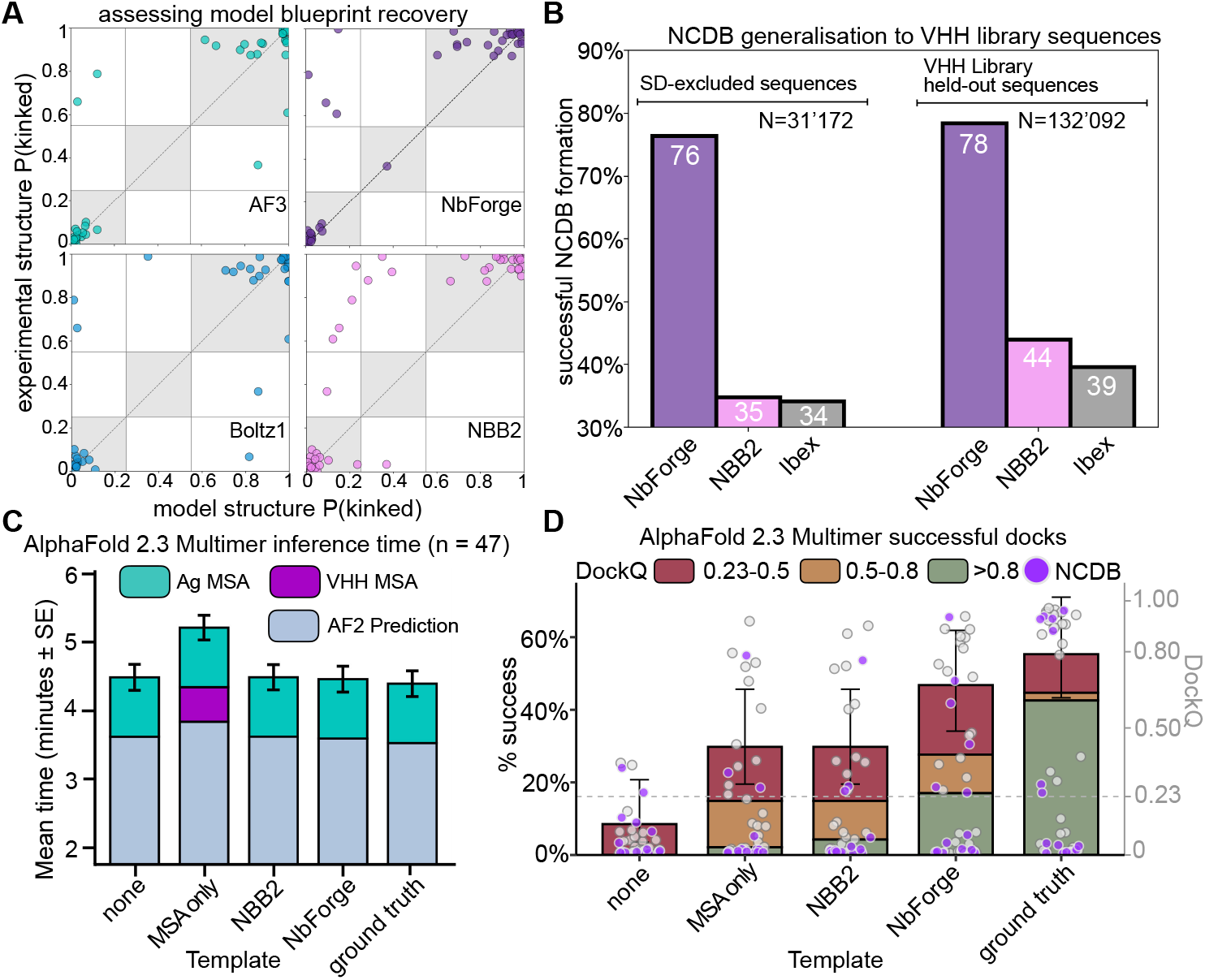
HCDR3 blueprint agreement, non-canonical disulphide (NCDB) recovery, and downstream docking performance. (**A**) Agreement between model-predicted and experimentally observed HCDR3 blueprint states, shown as structure-classifier probabilities P(kinked) for experimental structures (y-axis) versus predicted structures (x-axis) from AlphaFold3 (AF3), Boltz1, NbForge, and NanobodyBuilder2 (NBB2). Points represent individual nanobodies from the *n* = 47 test set (see also Table 3). (**B**) Bar plot reporting the fraction of sequences forming the expected NCDB for the lightweight models NbForge, NBB2, and Ibex on two different large sequence datasets; N indicates dataset size; values within bars indicate percentages. Both datasets contain only sequences with 4 or 5 cysteines, the two forming the canonical disulphide (AHo positions 23 and 106), one in the HCDR3 and another at either AHo positions 57 or 40. See Methods for more information. (**C**) and (**D**): AF2M inference time (C) and DockQ-based performance (D) in modelling *V*_*HH*_ -antigen complexes under different input conditions, including inputting monomeric templates predicted by NbForge or NBB2 (x-axis). In D, the left axis (bars) is the cumulative percent of structural models in each DockQ-accuracy group (legend), and the right axis (data points) is the DockQ score of each modelled structure. The “ground truth” group is obtained by providing to AF2M as template the monomeric nanobody structure as found in the experimentally determined bound complex, and it thus serves as a positive control for the highest achievable performance via templated predictions.

Despite this high performance, the remaining errors are informative. Across models, misclassifications are dominated by missed kinked blueprints (false negatives), i.e., predicting an extended blueprint when the experimental structure exhibits framework contacts characteristic of a kinked blueprint. This trend is most pronounced for NBB2, where some violations with the experimental structure’s kinked blueprint are recovered by the other models (Figure 3E, 4). These examples underscore that, while blueprint recovery is picked up by the models implicitly, the structure module on its own with only backbone coordinate loss terms (the main head of the NBB2 architecture) can still fail to capture subtle loop-framework packing patterns that the larger models recover more consistently.

### Non-canonical disulphide bridge recovery

Recovery of non-canonical disulphide bridges separates methods more strongly than kinked/extended blueprints (Table 3). AF3 recovers 14/15 (93.3%), NbForge recovers 13/15 (86.7%), and Boltz1 recovers 11/15 (73.3%), whereas NBB2 recovers only 4/15 (26.7% - exact McNemar p-value vs NbForge 0.004). The failure pattern in NBB2 is qualitatively consistent with a HCDR3 misplacement: many of the NCDB cases where NBB2 fails are ones that the other models recover (Figure 3F), and these same targets correspond to some of NbForge’s most accurate HCDR3 predictions (Table 3, 4), suggesting that failing to satisfy the covalent constraint often coincides with broader loop misplacement. At the other extreme, one NCDB target (Figure 3D) is missed by all models, and is also among the highest-error HCDR3 cases for the strongest predictors (PDB ID: 8zer). Notably, this NCDB corresponds to cysteines that are in an unorthodox position of typical NCDB cysteine positions in *V*_*HH*_ AHo sequence highlighted in Figure 1 (position 32 and 111 respectively).

**Table 4:** Median Backbone coordinate accuracy (RMSD, Å) stratified by presence of a non-canonical disulphide bond (NCDB *n* = 15) or by its absence (no NCDB *n* = 32). RMSDs are calculated for all backbone heavy atoms following framework superimposition and reported as median with bootstrap 95% confidence intervals.

| NCDB | Region | AF3 | Boltz1 | NBB2 | NbForge |
| --- | --- | --- | --- | --- | --- |
| no NCDB | FR | 0.70 [0.58, 0.73] | 0.66 [0.58, 0.74] | 0.76 [0.62, 0.87] | 0.73 [0.68, 0.83] |
|  | HCDR1 | 1.20 [0.94, 1.67] | 1.18 [0.96, 1.55] | 1.74 [1.31, 2.23] | 1.61 [1.15, 2.11] |
|  | HCDR2 | 0.68 [0.53, 1.06] | 0.74 [0.58, 1.42] | 0.95 [0.83, 1.43] | 1.05 [0.89, 1.33] |
|  | HCDR3 | 2.75 [1.59, 3.55] | 2.92 [2.11, 4.12] | 3.56 [2.91, 5.01] | 2.95 [2.51, 3.71] |
| NCDB | FR | 0.68 [0.52, 0.76] | 0.65 [0.54, 0.77] | 0.85 [0.67, 0.88] | 0.70 [0.64, 0.85] |
|  | HCDR1 | 1.19 [0.85, 1.97] | 1.63 [0.83, 2.73] | 1.72 [1.48, 2.62] | 1.65 [1.39, 1.90] |
|  | HCDR2 | 1.26 [0.70, 1.37] | 1.26 [0.63, 1.79] | 1.44 [1.08, 1.73] | 1.26 [1.00, 1.64] |
|  | HCDR3 | 2.97 [2.12, 3.70] | 3.58 [2.30, 3.74] | 3.45 [2.99, 4.53] | 2.95 [2.69, 3.53] |

Next, we stratified RMSD by NCDB status. As expected, the NCDB-bearing subset is enriched for substantially longer HCDR3s (mean 19.6 residues versus 13.9 residues in the non-NCDB subset), reflecting a different region of sequence and conformational space than most HCDR loops present in standard *V*_*HH*_ benchmarks. Despite the longer loop, NbForge maintains strong HCDR3 accuracy on NCDB-bearing targets and remains among the top-performing methods in this subgroup (median HCDR3 RMSD 2.95 Å for NbForge, compared with 2.97 Å for AF3, 3.58 Å for Boltz1, and 3.45 Å for NBB2; Table 4, Figure 3). Boltz1 exhibits a modest performance degradation in median HCDR3 RMSD on NCDB-bearing targets relative to the non-NCDB subset (2.92 *→* 3.58 Å). NBB2, unlike the other models, consistently performs worse on both classes of NCDB and no NCDB cases (3.45 and 3.74 Å respectively), whereas NbForge’s and AF3’s performances remain sub 3 Å between NCDB and no-NCDB structures (NbForge: 2.95 *→* 2.95 Å and AF3: 2.75 *→* 2.97 Å). This pattern is consistent with the intended role of the explicit disulphide-aware objective, where within a covalently constrained loop, correctly recovering the disulphide bridge can reduce the effective loop conformational space and stabilize loop placement (Figure 3C, D, F).

### Inference time

Prediction time is a key feature of folding models, as faster inference time can enable large-scale dataset evaluation of candidates from design campaigns or laboratory library-screening without incurring prohibitive compute costs. We measured end-to-end wall-clock inference time on the 47-target test set (Table 5). While AlphaFold3 and Boltz1 require on the order of tens of seconds per structure on GPU (∼ 1 minute with MSA, ∼ 40 s without MSA), NbForge predicts structures in less than a second per sequence on both CPU and GPU (Table 5). This throughput difference, combined with NbForge’s competitive HCDR3 RMSD, strong blueprint recovery and NCDB formation, is decisive to enable screening-scale *V*_*HH*_ structure prediction in discovery workflows where thousands to millions of candidates may be triaged.

**Table 5:** Wall-clock inference timings on the NbForge test-set (*n* = 47) are reported in seconds and averaged over three independent runs on an NVIDIA Quadro RTX 8000 GPU and an Intel Xeon Platinum 8263. ^†^ For AlphaFold3 and Boltz1, values in parentheses correspond to predictions performed without MSA/template search. ^‡^ For NbForge, values in parentheses correspond to the extra time taken by the optional OpenMM relaxation step, with monomer inference times preceding. NBB2 timings are CPU-only.

| Metric | AF3 | Boltz1 | NBB2 | NbForge-GPU | NbForge-CPU |
| --- | --- | --- | --- | --- | --- |
| Total time (s; 3 runs)<sup>†‡</sup> | 2676.9 (1841.8) | 2961.4 (2088.1) | 410 | 6.1 (72.2) | 10.0 (128.1) |
| Per-structure time (s)<sup>†‡</sup> | 57.0 (39.2) | 63.0 (44.4) | 4.5 | 0.13 (1.54) | 0.21 (2.73) |

### Implications for nanobody design and discovery

In library triage or de novo nanobody generation, misplacing a disulphide-constrained HCDR3 can yield a fold that appears geometrically plausible yet is fundamentally incompatible with the relevant paratope conformation. To address whether NbForge can correctly predict NCDBs at library scale, we compared NbForge against two lightweight models that offer comparable speed: NBB2 and Ibex, a recently introduced nanobody predictor trained on public structural data [48]. We predicted structures for two stringent sequence sets that were excluded from our main training and self-distillation pipelines: (i) self-distillation-excluded sequences and (ii) *V*_*HH*_ -library held-out sequences (see Methods). To increase confidence that sequences in these sets can in principle form an NCDB, we restricted the analysis to sequences containing four or five cysteines, including a cysteine at AHo position 40 or 57, which are the dominant cysteine positions observed in four-/five-cysteine nanobody structures (Figure 1). Because Ibex’s training data overlap with our curated NbForge test set, we did not include Ibex in the test-set benchmark; however, Ibex provides a useful comparator for the library-scale NCDB screen where the analysed sequences are distinct from both training and test sets. Under this protocol, NbForge recovered NCDB conformations far more frequently than the other models across both datasets, forming NCDBs in 76.4% (self-distillation-excluded) and 78.4% (*V*_*HH*_ -library held-out) of sequences, compared with 34.8%/44.1% for NBB2 and 34.1%/39.4% for Ibex (Figure 4B). While NbForge significantly improves in the HCDR3-NCDB recovery of these sequences relative to the other lightweight models, which we show corresponds to lower HCDR3-RMSD, these lightweight predictors are far from achieving complete NCDB recovery. We also note that sequence-based filtering cannot guarantee that every multi-cysteine nanobody sequence from camelid *V*_*HH*_ repertoires will in fact adopt a NCDB architecture, and we optimize for this by selecting the most likely of cysteine-pairing partners. Yet, higher NCDB recovery at screening scale implies fewer false negatives when prioritizing candidates with constrained HCDR3 architectures, thereby expanding the set of sequences that can be advanced for experimental validation rather than being discarded due to modelling failures in disulphide-constrained loops.

Beyond improving backbone structure accuracy, blueprint, and NCDB recovery, NbForge yields tangible gains in downstream nanobody-antigen modelling. Using AlphaFold 2.3 Multimer on the NbForge test-set of bound complexes, which unlike AF3 is an open-source model, we compared runs using (i) no nanobody-side evolutionary/template information, (ii) *V*_*HH*_ MSA-only, (iii) NBB2 monomer templates, (iv) NbForge monomer templates, and (v) the experimental monomer from the bound complex as a positive-control template (Figure 4C–D). NbForge templating increased the fraction of successful complex predictions (DockQ 0.23) to ∼ 47%, compared with ∼ 30% for *V*_*HH*_ MSA-only or NBB2 templating and ∼ 10% with no nanobody-template input (Figure 4D). In addition, NbForge shifted the DockQ distribution toward higher-quality predictions, increasing the proportion of high-accuracy models (DockQ > 0.8) relative to MSA-only/NBB2 and approaching the ceiling set by the experimental-template control (Figure 4D). These gains are achieved without increasing AF2M wall-clock time relative to non-MSA baselines; the main runtime penalty arises from *V*_*HH*_ MSA generation in the MSA-only condition (Figure 4C). Finally, several NCDB-bearing nanobodies (purple points) that fail under MSA-only inputs are rescued by NbForge templating, indicating that enforcing nanobody-specific inductive biases can improve complex prediction even for disulphide-constrained HCDR3 loops (Figure 4D). Together, these results suggest that explicitly enforcing nanobody-feature inductive biases during training can translate directly into more reliable complex prediction, strengthening the utility of NbForge as a scalable model for discovery pipelines that prioritize rapid screening, epitope mapping, and structure-guided optimization.

## 3 Discussion

Accurate structural modelling of nanobodies (*V*_*HH*_ s) is becoming increasingly valuable for nanobody discovery and engineering efforts. However, the regions that typically determine binding and developability potential are also those that remain more difficult to model. HCDR3 loops span broad conformational diversity, and many nanobodies contain additional cysteines that form non-canonical disulphide bridges (NCDBs) that impose covalent constraints on loop geometry. Standard benchmarking of structure-prediction tools focuses on continuous coordinate errors (typically RMSD), yet two models can have similar RMSD while differing in discrete structural commitments that are directly relevant to paratope architecture – most notably the global HCDR3 blueprint (kinked versus extended) and the disulphide connectivity. Taken together, our results support reporting HCDR3-blueprint and disulphide-connectivity recovery alongside coordinate accuracy when assessing nanobody folding models, and demonstrate that incorporating these features as inductive biases improves *V*_*HH*_ structure prediction, particularly for long, cysteine-rich HCDR3 loops.

This work introduces two components that make these discrete structural commitments explicit. NbFrame provides a sequence-level prior over kinked and extended HCDR3 regimes using hallmarks concentrated at the CDR3–framework interface, allowing users to flag predictions whose loop placement is inconsistent with the expected blueprint [49, 50, 31]. Practically, such hallmarks also provide a compact set of positions to preserve when proposing framework mutations for humanisation or developability engineering, because altering them can shift the loop into a different blueprint state even when the CDR sequences are unchanged. NbForge encodes these commitments both at the objective level, via an explicit loss term encouraging physically valid disulphide geometry, and at the data level, via self-distillation on a training set filtered to retain only structures that satisfy disulphide bridge formation and the NbFrame blueprint prior. Together, these signals improve HCDR3 RMSD and increase HCDR3 blueprint and NCDB recovery relative to NBB2, while reaching accuracy at par with much deeper models such as AF3 and Boltz (Table 3, 4, Figure 3).

Discovery and engineering pipelines increasingly require both accuracy and throughput. Lightweight predictors such as NBB2 have already been deployed in functional optimisation and developability-focused studies of *V*_*HH*_ s [51, 52, 53, 54]. NbForge was designed to deliver backbone accuracy and feature recovery without the latency of large, MSA-dependent deep architectures, enabling repertoire-scale inference where structure prediction becomes a routine filtering step (Figure 4D). Using NbForge monomers as templates for AlphaFold-Multimer increases the fraction of successful complexes (DockQ *≥* 0.23) and shifts predictions toward higher-quality outcomes (DockQ *≥* 0.8) relative to NBB2 templates or the default implementation (Figure 4B, D). Given that AlphaFold-Multimer has been central to multiple design campaigns that yielded successful nanobody designs [55, 51], these results support the practical importance of reliable monomer geometry for downstream complex prediction.

High-quality coordinate and surface representations are also important for residue-burial analysis, and surface-level engineering tasks that are sensitive to loop placement and covalent topology [51, 52, 46, 56], including nanobody humanization via framework resurfacing [52] and structure-guided affinity maturation [57]. Moreover, Bashour *et al*. showed that developability metrics derived from predicted antibody structures are captured more reliably when dynamics are incorporated via molecular dynamics simulations, particularly for properties driven by surface exposure, charge redistribution, and loop flexibility [21]. In this context, backbone accuracy and feature recovery are essential, as simulations initiated from structures with incorrect covalent topology or unrealistic heavy-atom geometries (most notably the cysteine side-chain geometry required for disulphide formation, and the short-range heavy-atom contacts that define the blueprint states) [35] fail to explore physically meaningful conformational sub-states, resulting in misleading flexibility and developability assessments.

NCDBs are common in camelid repertoires and often act as stabilizing topological constraints, while also introducing additional folding and redox sensitivities [58]. Consistent with this dual role, mutagenesis and structural studies show that removing NCDBs can reduce thermal stability and shift reversible unfolding toward aggregation-prone behaviour, and in some cases reduce binding affinity [34, 59]. In *de novo V*_*HH*_ design, introducing additional cysteines into CDRs is typically avoided to limit combinatorial complexity and manage developability risk associated with redox sensitivity. Our results suggest that, with improved disulphide-connectivity recovery, this design constraint can be relaxed when the goal is to explore disulphide-stabilized paratopes. NbForge allows modelling of NCDB-bearing HCDR3s, providing an explicit mechanism to pre-organize paratopes to achieve binding modes inaccessible without NCDBs, and to reduce conformational heterogeneity, likely limiting unfavourable entropy costs upon binding [60]. Moreover, the non-canonical cysteine-pair positional information (Figure 1D, S7) provides concrete guidance for selecting sequence sites when engineering disulphide-stabilized loops in design campaigns.

While accurate monomeric *V*_*HH*_ structure prediction and feature recovery helps produce backbones closer to experimentally observed structures, proteins populate a dynamic equilibrium of different interconverting conformational states in solution [60]. Indeed, recent works have focused on developing and using deep-learning methods to predict protein thermodynamic ensembles from sequence [61, 62, 63]. Our results suggest that such predicted ensembles should be benchmarked beyond coordinate uncertainty and dispersion alone, where dispersion can be high either because the model samples alternative but topologically consistent states, or because it produces chemically invalid conformers. For example, ensemble-level metrics such as the fraction of samples satisfying disulphide constraints and the distribution of NbFrame blueprint scores may help distinguish physically plausible heterogeneity from sampling artifacts, and better support downstream interpretations and simulations.

Finally, the ground truth experimental structures in the PDB are a crystallographic or Cryo-EM snapshot, most of which captured in the antigen-bound state [64]. Because our held-out benchmark consists of *V*_*HH*_ s taken from bound complexes, our performance metrics quantify agreement with bound conformations rather than the unbound solution ensemble. Antigen binding may shift CDR conformational populations (most prominently HCDR3) consistent with conformational selection, and in some cases additional induced-fit rearrangements are observed [65, 66]. However, a recent analysis of bound and unbound antibody PDB structures suggests that this behaviour is rare, as these two conformations are typically very similar [67]. In general, a predictor that accurately represents the unbound ensemble may be penalized when scored against a bound reference, while a predictor biased toward bound-like geometries could appear to dock better despite being less representative of the *V*_*HH*_ free state in solution. This disparity motivates future investigations on paired bound/unbound structures and ensemble-aware reference representations (e.g., MD-derived distributions), to ultimately provide more reliable foundations for downstream dynamics-based developability and functional analyses [60, 21].

We release NbForge and NbFrame as open-source, downloadable software and we make available a user-friendly web server for NbForge structural modelling. We intend these as community resources for routine, high-throughput nanobody modelling, where rapid access to reliable unbound structures enables earlier and broader use of structure-informed decision making.

## 4 Methods

### 4.1 NbFrame: HCDR3 conformation classification

#### Structural database curation for NbFrame training

A total of 1,056 *V*_*HH*_ structures were initially retrieved from SAbDab databse [37]. Structures were superimposed to a reference nanobody (PDB: 2P49, chain B) using framework residues only and a quality filter excluded structures with framework RMSD > 2.0 Å to the reference, removing 51 poorly resolved or domain-swapped structures and yielding 1,005 structures. To prevent data leakage from redundant sequences, we performed CDR-based deduplication. Structures sharing identical HCDR1+HCDR2+HCDR3 sequences (concatenated as a single string) were identified, and one representative per unique combination was retained, yielding a final dataset of 929 unique *V*_*HH*_ structures. The representative structure was chosen based on minimal number of missing residues (i.e. without assigned coordinates) in the core region (i.e. excluding 4 residues at N- and C-terminus) and highest resolutions. Sequences were aligned and numbered according to the AHo numbering scheme [68], providing a consistent 149-position alignment across structures. HCDR boundaries were defined as: HCDR1 (AHo 27–42), CDR2 (AHo 57–69), and HCDR3 (AHo 108–138).

A temporal split based on PDB accession codes was used for NbFrame only, where PDB structures with ID’s beginning with “9” were designated as the held-out NbFrame test set (*n* = 100) (Figure S10), while remaining structures formed the NbFrame training set.

#### NbFrame Structural feature calculation

Six features were calculated for each structure to characterize HCDR3 conformation:

#### Backbone angles

The *α* (virtual bond angle) and *τ* (virtual torsion angle) were calculated at both the N-terminal and C-terminal stems of the HCDR3 loop. For the N-terminal stem, angles were measured at AHo positions 105–108; for the C-terminal stem, at positions 135–138. Virtual bond angles were defined as:

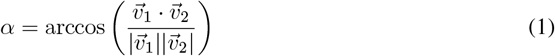

where 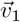 and 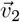 are vectors between consecutive C*α* atoms.

#### Contact density

Atomic contacts between HCDR3 residues and FR2 residues (AHo positions 44–55) were counted using a distance cut-off of 4.5 Å. CDR3 stem residues (AHo positions 108, 109, 136, 137, 138) were excluded from contact counting. Contact density was normalized by the full CDR3 length to provide a length-independent measure:

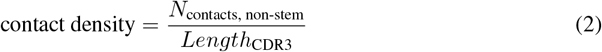

where *N*_contacts, non-stem_ is the number of contacts between non-stem CDR3 residues and FR2, and *Length*_CDR3_ is the full CDR3 length.

#### FR2 relative solvent accessibility

Solvent accessible surface area (SASA) was calculated using FreeSASA [69] with default parameters (probe radius 1.4 Å). Relative solvent accessible surface area (RSASA) was computed for key FR2 positions (44 and 54) by normalizing with theoretical maximum SASA values derived from exhaustive conformational sampling of Gly-X-Gly tripeptides [70]. The feature fr2_rsa_key was defined as the mean RSA of AHo positions 44 and 54.

#### Unsupervised clustering and manual labelling of kinked and extended blueprints

To identify natural groupings in the six-dimensional feature space, we applied Gaussian Mixture Model (GMM) clustering with full covariance matrices using scikit-learn [71]. Features were standardized using StandardScaler prior to clustering. Model selection based on silhouette score indicated an optimal two-cluster solution (silhouette = 0.34), consistent with the hypothesized kinked/extended dichotomy.

For ground truth establishment, we sampled 50 structures from each GMM cluster, including: i) structures from each cluster core (GMM probability > 0.99), (ii) structures from near-boundary regions (probability 0.90–0.99), and (iii) structures from the true boundary (probability 0.10–0.90). Each structure was manually labelled as Kinked (K) or Extended (E) based on visual assessment of HCDR3 conformation. Labelling criteria were kinked: HCDR3 bends back toward the framework and makes contacts with FR2 residues; Extended: HCDR3 projects outward with minimal or no FR2 contact.

Manual classification yielded 60 extended and 40 kinked labels from the 100 sampled structures, reflecting 80% concordance between GMM clustering and manual labels. Analysis of the 20 discordant cases revealed that GMM clustering primarily captured C-terminal backbone geometry (particularly *α*_*C*_ angles), whereas the structural-biology definition of kinked conformations requires not only kinked-like angles but also physical CDR3-FR2 contacts that shield hydrophobic framework residues from solvent. Among the discordant structures, 15 exhibited kinked-like angles but lacked FR2 contacts (contact density = 0) and were therefore classified as extended, while only 5 structures from the extended cluster were reclassified as kinked upon visual inspection.

#### NbFrame structure classifier training

A logistic regression classifier was trained on the 100 manually labelled structures, using stratified 5-fold cross-validation. Features were standardized using StandardScaler, and the scaler and classifier were combined into a single pipeline. Model performance was evaluated using cross-validation on the training set (ROC-AUC = 0.96) and then on the 100 held-out test structures with manual labels (ROC-AUC = 0.99, accuracy = 94%).

The NbFrame structural classifier computes the probability of a kinked conformation as:

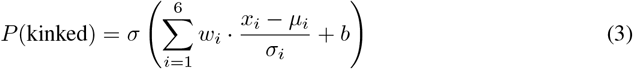

where *σ*(*z*) = 1*/*(1 + *e*^*−z*^) is the sigmoid function, *x*_*i*_ is the raw feature value, *µ*_*i*_ and *σ*_*i*_ are the per-feature mean and standard deviation from the training set (used for standardization), *w*_*i*_ is the learned coefficient for feature *i*, and *b* = *−*0.352 is the intercept term. The complete parametrisation is provided in Table 6.

**Table 6:** NbFrame structure classifier parameters. Features are standardized by subtracting the mean (*µ*) and dividing by the standard deviation (*σ*) before applying the logistic regression coefficients (*w*). Training set: *n* = 100 manually labelled structures.

| Feature | Mean ($\mu$) | Std ($\sigma$) | Coefficient ($w$) |
| --- | --- | --- | --- |
| $\alpha_N$ (degrees) | -62.09 | 133.53 | +0.11 |
| $\tau_N$ (degrees) | 126.22 | 11.47 | +0.60 |
| $\alpha_C$ (degrees) | -13.31 | 101.63 | +0.48 |
| $\tau_C$ (degrees) | 109.68 | 13.78 | -0.42 |
| contact density | 0.29 | 0.31 | +1.86 |
| fr2_rsa_key | 0.14 | 0.09 | -1.16 |
| Intercept: $b = -0.352$ | | | |

**Table 7:** Sequence hallmark features used for the NbFrame sequence classifier, derived from 829 unique *V*_*HH*_ sequences from SAbDab [37], with blueprint labels assigned by the structure classifier. Features represent specific (AHo position, amino acid) combinations with corresponding enrichment in Kinked versus Extended structures. Log_2_FC indicates Fold-Change of frequencies; positive values indicate enrichment in Kinked structures. Frequencies are provided as percentages from a total of 478 kinked and 351 extended structures, and the p-values is from a Fisher’s exact test.

| Region | AHo Position | AA | $\text{Log}_2\text{FC}$ | p-value | % Kinked | % Extended |
| --- | --- | --- | --- | --- | --- | --- |
| *Framework 1* |  |  |  |  |  |  |
| FR1 | 12 | S | +1.14 | $5.4 \times 10^{-5}$ | 17.6 | 8.0 |
| FR1 | 12 | L | -0.14 | $2.4 \times 10^{-3}$ | 78.7 | 86.9 |
| FR1 | 15 | P | -0.58 | $3.9 \times 10^{-6}$ | 32.2 | 48.1 |
| FR1 | 15 | A | +0.38 | $3.9 \times 10^{-5}$ | 62.3 | 47.9 |
| FR1 | 17 | D | +2.32 | $8.5 \times 10^{-5}$ | 7.1 | 1.4 |
| *Framework 2* |  |  |  |  |  |  |
| FR2 | 44 | F | +1.72 | $< 10^{-45}$ | 81.0 | 24.5 |
| FR2 | 44 | Y | -2.77 | $< 10^{-45}$ | 9.0 | 61.3 |
| FR2 | 51 | E | +0.72 | $1.2 \times 10^{-20}$ | 79.1 | 47.9 |
| FR2 | 51 | Q | -2.63 | $1.7 \times 10^{-25}$ | 5.2 | 32.5 |
| FR2 | 54 | F | +1.24 | $2.1 \times 10^{-15}$ | 45.2 | 19.1 |
| FR2 | 54 | G | +2.21 | $9.0 \times 10^{-18}$ | 29.1 | 6.3 |
| FR2 | 54 | L | -2.34 | $1.8 \times 10^{-26}$ | 7.3 | 37.0 |
| FR2 | 54 | W | -1.00 | $6.5 \times 10^{-6}$ | 11.9 | 23.9 |
| FR2 | 56 | A | -0.20 | $1.7 \times 10^{-3}$ | 68.0 | 78.1 |
| *Framework 3* |  |  |  |  |  |  |
| FR3 | 85 | S | -1.73 | $3.7 \times 10^{-7}$ | 4.4 | 14.5 |
| FR3 | 85 | A | +0.21 | $5.9 \times 10^{-5}$ | 84.3 | 72.6 |
| FR3 | 103 | V | -0.29 | $5.4 \times 10^{-7}$ | 68.0 | 83.2 |
| FR3 | 103 | I | +1.45 | $1.3 \times 10^{-4}$ | 11.7 | 4.3 |
| FR3 | 107 | A | +1.20 | $< 10^{-45}$ | 86.8 | 37.9 |
| FR3 | 107 | N | -2.46 | $8.3 \times 10^{-21}$ | 5.2 | 28.8 |

For deployment, confidence thresholds were implemented: predictions with *P* (kinked) > 0.7 were labelled as “kinked,” *P* (kinked) < 0.3 as “extended,” and intermediate probabilities as “unclear.” The trained classifier was applied to all 929 structures, yielding 480 Kinked (51.7%), 356 Extended (38.3%), and 93 Unclear (10.0%) predictions.

#### NbFrame sequence classifier

To enable sequence-based predictions, we developed a logistic regression classifier based on framework sequence features associated with HCDR3 blueprint. For each (AHo position, amino acid) combination in framework regions (in AHo numbering - FR1: positions 1–26; FR2: positions 43–56; FR3: positions 70–107; FR4: positions 139–149), we computed the log_2_ fold-change of frequencies (*Log*_2_*FC*) between kinked and extended structures:

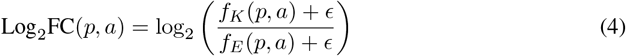

where *f*_*K*_(*p, a*) and *f*_*E*_(*p, a*) are the frequencies of amino acid *a* at AHo position *p* in Kinked and Extended structures, respectively, and *ϵ* = 0.5*/N* is a pseudo-count to avoid division by zero (*N* is the number of training structures).

For downstream analysis, we retained only those (*p, a*) combinations that satisfied: (i) Fisher’s exact test *p* < 0.05, and (ii) total observations *≥* 30 (across Kinked and Extended classes). For each combination, we computed: (i) point-biserial correlation with binary class labels in the training set structures (discriminative power), and (ii) Pearson correlation with structure classifier probabilities of training-set structures (consistency with structural predictions). The combined score was defined as the product of absolute correlations (|*r*_label_| *×* |*r*_struct_|), and the top 20 candidates by this metric were selected as hallmarks, because using 20 features maximised the *E/K* discrimination performance on the training set. This approach prioritizes features that are both discriminative and aligned with the structure classifier, such that the sequence classifier approximates structure-based classification from sequence alone. The resulting hallmarks are concentrated at biologically relevant positions: FR1 (positions 12, 15, 17), FR2 (positions 44, 51, 54, 56), and FR3 (positions 85, 103, 107). Features from CDR regions were excluded to avoid potential bias from CDR sequences in engineered or synthetic *V*_*HH*_ s.

A logistic regression model was trained on 829 sequences with known structures using soft labels from the structure classifier (*P* (kinked)) treated as continuous probability. For a given AHo-aligned sequence *S*, features are encoded as follows: for each hallmark *j ∈ {*1, …, 20*}*, defined by position *p*_*j*_ and amino acid *a*_*j*_ with associated Log_2_FC_*j*_ value (Table 7), the feature *x*_*j*_ is set to Log_2_FC_*j*_ if the sequence has amino acid *a*_*j*_ at position *p*_*j*_, and 0 otherwise:

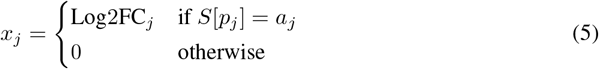

The probability of a kinked conformation is then computed as:

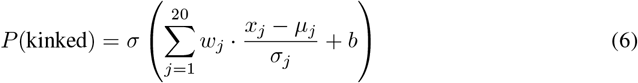

where *σ*(*𝓏*) = 1*/*(1 + *e*^*−𝓏*^) is the sigmoid function, *µ*_*j*_ and *σ*_*j*_ the mean and standard deviation of feature *x*_*j*_ computed from the training set, and *w*_*j*_ and *b* are the learned model coefficients and bias, respectively.

The model was evaluated on the 100 held-out structures with manual labels as ground truth, achieving ROC-AUC of 0.94 and accuracy of 86%.

#### NbFrame software implementation

NbFrame is implemented in Python and is available as an open-source package on GitHub (https://github.com/Mateusz-Jaskolowski/NbFrame). The structure classifier accepts PDB or mmCIF files and outputs predicted class and probability. The sequence classifier accepts amino acid sequences (with optional automatic AHo alignment) and outputs probability of kinked or extended.

### 4.2 NbForge Model Architecture

#### Disulphide bridge loss

We adapt the architecture and base training objectives of ABody-Builder3 [72, 73], which in turn are built on the structure module of AlphaFold2. We augment these with a composite disulphide-bridge loss to encourage physically realistic cysteine pairing and geometry based on ground-truth disulphide bridges of *V*_*HH*_ s. All disulphide-specific loss terms are applied only to cysteine residues and are annealed during training.

Distance-based disulphide terms use linear hinge penalties; the S–S–C*β* bond angle is regularized with a Huber loss, while the *χ*_3_ dihedral is supervised using a bimodal wrapped likelihood.

For a nanobody containing *N ∈ {*2, 3, 4, 5*}* cysteine residues, let **s**_*i*_ *∈* ℝ^3^ and **c**_*i*_ *∈* ℝ^3^ denote the predicted SG and C*β* atom coordinates of cysteine *i*, respectively, and let 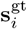 denote the corresponding ground-truth SG coordinate. All distances are measured in Å.

#### Cysteine pairing

To identify disulphide bonds, we compute the ground-truth SG–SG distances

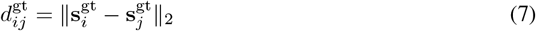

for all cysteine pairs using *i < j* to prevent double counting. Pairs satisfying

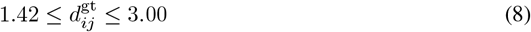

are considered candidates based on the distribution of SG-SG distances in ground-truth structures. From these, we greedily select a non-overlapping set of bonded cysteine pairs ℬ, such that each cysteine participates in at most one bond. All remaining cysteine pairs form the complementary set ℬ′.

#### SG–SG distance band and non-bonded repulsion

Let

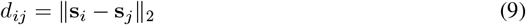

denote the predicted SG–SG distance. We apply a hinge-style penalty that constrains bonded cysteine pairs to a narrow distance band [1.84, 2.26] Å and repels additional cysteines to prevent unphysical “clusters” of cysteine residues “attracting” each others (by pushing any upaired cysteine to have their distance at least 4.1 Å away from the paired cysteines):

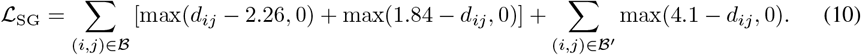

#### C*β*–C*β* distance prior

For bonded cysteine pairs, we regularize the predicted C*β*–C*β* distance using a flat-bottom hinge loss that penalizes deviations outside a target interval [2.85, 4.50] Å

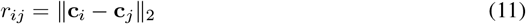

using a flat-bottom hinge loss:

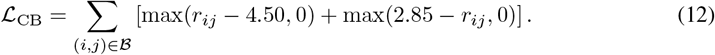

#### SG–SG–C*β* bond angle prior

For each bonded cysteine pair (*i, j*), we compute the two S–S–C*β* angles

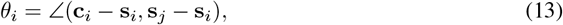

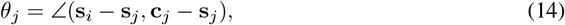

and penalize deviations from a target angle *θ* = 2.0 rad using a Huber loss with transition parameter *δ* = 0.04:

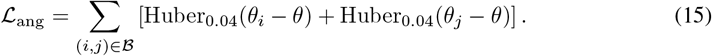

#### disulphide *χ*_3_ dihedral prior

For each bonded cysteine pair, we define the disulphide *χ*_3_ dihedral angle

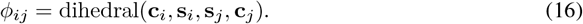

We impose a bimodal wrapped Gaussian penalty corresponding to the two canonical disulphide rotamers, providing a smooth objective that avoids hard mode assignment:

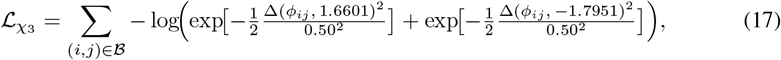

where Δ(*·, ·*) denotes the angular difference wrapped to (*−π, π*].

#### disulphide distogram loss

In addition to coordinate-based penalties, we supervise the predicted residue–residue distogram for bonded cysteine pairs only. Let 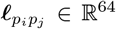 *∈* ℝ^64^ denote the predicted distogram logits between the sequence positions *p*_*i*_ and *p*_*j*_. We define a target bin corresponding to a SG–SG distance of 2.05 Å and apply a symmetric cross-entropy loss:

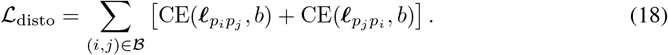

#### Annealing

The total disulphide objective is given by

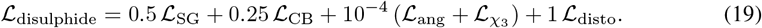

and is multiplied by an annealing factor that increases linearly from 0 to 1 over the first 100000 training steps.

#### Training and inference

A first version of the model was trained only on the *V*_*H*_ /*V*_*HH*_ experimental structures for 500,000 steps. This model was used to create the self-distillation set (Figure 2). Then, another version of the model was trained only on the filtered self-distillation dataset always for 500,000 steps. This model was then fine-tuned on the experimental *V*_*H*_ /*V*_*HH*_ structures for 500,000 steps. When performing inference, unless stated otherwise, structures predicted by NbForge were subjected to restrained energy minimisation in OpenMM using the AMBER14SB protein force field, as done by NBB2 [45, 74, 75], including the application of harmonic positional restraints to N, C*α*, C and C*β* atoms to keep the refined backbone close to its original prediction.

### 4.3 Evaluation of blueprint and NCDB features

#### Blueprint recovery

The NbFrame structure classifier was used to evaluate all ground truth, experimental structures of the test set. The classifier threshold of less than 0.25 is used to classify HCDR3 loop to be extended, in between 0.25 and 0.55 are deemed to be unclear, and greater than 0.55 is classified as kinked, as per the accuracy analysis performed in Figure S11. We then evaluated the predicted structures from each model using the same NbFrame structure classifier, and the overlap into the three different bins is used to measure the accuracy of each models’ structure relative to the experimental structures (Figure 4B, Table 3).

#### NCDB recovery

Both canonical and non-canonical disulphide bridges from all the SAbDab experimental *V*_*HH*_ s are evaluated. Only SG-SG distances of less than 3 Å were kept for evaluation based on previous work that used a similar distance threshold [76]. From this set of disulphide bridges, the 99th percentile of SG-SG distances was used (2.46 Å) as a distance threshold cutoff for the evaluation. After prediction, the expected disulphide-bridges and their respective cysteine SG atom distances were calculated and if they satisfied the 2.46 Å threshold they were deemed to have formed the disulphide bridge. The violations of this 2.46 Å SG-SG distance threshold for each predicted structures are reported in Table 3.

### 4.4 NCDB recovery in large sequence datasets

Two large sequence datasets were folded with the three lightweight models (NbForge, NBB2, and Ibex) and evaluated for their NCDB recovery (Figure 4). Dataset 1 (self distillation-excluded sequences) was derived from an initial set of 154,115 sequences for which structures were predicted with the teacher *V*_*H*_ */V*_*HH*_ model. After filtering out structures for structural violations (Figure 2D, see Results), 95,540 sequences remained. From these, we retained only sequences with (i) 4–5 cysteines total across the full *V*_*HH*_ domain sequence, (ii) a cysteine at AHo position 40 and/or 57 (dominant cysteine position shown in Figure 1B), and (iii) at least one cysteine in HCDR3 (i.e., enabling a putative 40/57–HCDR3 NCDB), yielding 31,172 sequences (SD-excluded dataset). Dataset 2 (held-out *V*_*HH*_ library) started from sequences retained after the “cluster by 80% MMSEQ” and “remove singletons” steps (Figure 2D). We removed any sequences overlapping Dataset 1 (exact sequence identity) and then applied the same cysteine-based criteria as done for Dataset 1, yielding 132,103 sequences.

### 4.5 AlphaFold 2.3 Multimer *V*_*HH*_ -antigen complex prediction

All nanobodies from the NbForge test-set are in the bound state against their respective antigen in their PDBs. Multi sequence alignment (MSAs) were generated separately for both the antigen and the nanobody against the UniRef30 clustered database and the ColabFold environmental sequence database using MMseqs2 [42]. Structure prediction was performed using ColabFold running AlphaFold 2.3 Multimer (AF2M) [13, 77, 78], with custom monomeric *V*_*HH*_ templates either from experimental structures, NBB2, or NbForge modelled structures, with the precomputed antigen MSAs from above supplied as input. Predictions used the default five model-prediction [78] with three recycle iterations and a single random seed (0). DockQ value was calculated on the whole nanobody-antigen complex, and the top DockQ score was selected from the 5 replicates of each complex prediction respectively [79].

### 4.6 Heavy-model monomer inference pipelines

AlphaFold3 (AF3) structure prediction was performed in single-chain (monomer) mode for all *V*_*HH*_ sequences analysed in this work. For each sequence, a multiple sequence alignment (MSA) was generated using a local MMseqs2-based ColabFold-style pipeline [77], producing an.a3m alignment that was provided to AF3 as a custom MSA input. Structural templates were not provided manually; instead, AF3 was run with template usage enabled and performed automatic template selection via its internal template search. AF3 inference was run using a single model seed; this seed produces multiple diffusion samples (five by default), from which the highest-ranked model under AF3’s default ranking/scoring scheme was retained as the single structural prediction for each sequence. This top-ranked model was used for all downstream analyses and visualisation.

Boltz1 structures were predicted with the official Boltz implementation using the Boltz-1 model weights. Each target sequence was provided as a single-chain input (*V*_*HH*_ domain only), and we generated one predicted structure per target (one diffusion sample) for downstream evaluation, consistent with the default (`–diffusion_samples 1`). Inference used the default Boltz prediction hyperparameters: 3 recycling steps (`–recycling_steps 3`) and 200 sampling steps (`–sampling_steps 200`). Multiple sequence alignments (MSAs) were used for the primary Boltz1 predictions. MSAs were generated using the standard Boltz workflow via the MMseqs2 MSA server (`–use_msa_server`), which queries the ColabFold MMseqs2 API endpoint by default. For monomeric inputs, a single.a3m MSA is auto-generated when `–use_msa_server` is enabled. No structural templates were provided for Boltz1 predictions, which (unlike AlphaFold3) cannot use input templates in its inference pipeline.

## 5 Code and data availability

The NbFrame source code, sequence and structure models and weights are available at https://github.com/Mateusz-Jaskolowski/NbFrame. The NbForge source code and weights are available at https://gitlab.doc.ic.ac.uk/sormanni-lab/nbforge. All the parsed training, validation and testing datasets employed in this study are available alongside the source code.

## 6 Author Contributions

MA and PS conceived the project. MA conducted all formal analysis and methodology development of NbForge. MJ conducted all formal analysis and methodology of NbFrame. MA and MJ incorporated and analysed NbFrame in the context of NbForge. All authors contributed to the development of the pipelines and analysis. MA and PS wrote the first version of the paper; all authors edited the paper.

## 7 Acknowledgments

PS is a Royal Society University Research Fellow (grant no. URF\R\251013). We acknowledge funding from UK Research and Innovation (UKRI) Engineering and Physical Sciences Research Council (EPSRC grant no. EP/X024733/1, an ERC starting grant to PS underwritten by UKRI) and Biotechnology and Biological Sciences Research Council (BBSRC BB/Y007816/1). MJ was supported by an SNSF Postdoc.Mobility Fellowship (grant no. P500PB_230458) funded by the Swiss National Science Foundation.

This work was supported by grants from the Norwegian Cancer Society Grant (215817, to VG), Research Council of Norway projects (300740, 331890 to VG). This project has received funding (to VG) from the Innovative Medicines Initiative 2 Joint Undertaking under grant agreement No 101007799 (Inno4Vac). This Joint Undertaking receives support from the European Union’s Horizon 2020 research and innovation programme and EFPIA. This communication reflects the author’s view and neither IMI nor the European Union, EFPIA, or any Associated Partners are responsible for any use that may be made of the information contained therein. Funded by the European Union (ERC, AB-AG-INTERACT, 101125630, to VG).

This work was supported in part by the Intramural Research Program of the National Institutes of Health (NIH) and the Division of Intramural Research (DIR), NIAID/NIH (THN, EG.). This study used the computational resources of the NIH High Performance Computing (HPC) Biowulf cluster (http://hpc.nih.gov) and the Office of Cyber Infrastructure and Computational Biology (OCICB) HPC Skyline cluster at NIAID/NIH, Bethesda, MD (https://skyline.niaid.nih.gov/). The contributions of the NIH authors are considered Works of the United States Government. The findings and conclusions presented in this paper are those of the authors and do not necessarily reflect the views of the NIH or the U.S. Department of Health and Human Services.

MA is supported by a Harding Distinguished Postgraduate Scholarship.

## 8 Disclosure statement

V.G. declares advisory board positions in aiNET GmbH, Enpicom B.V, Absci, Omniscope, and Diagonal Therapeutics. All other authors declare no potential conflict of interest.

## 9 Supplementary Information

**Figure S1:**
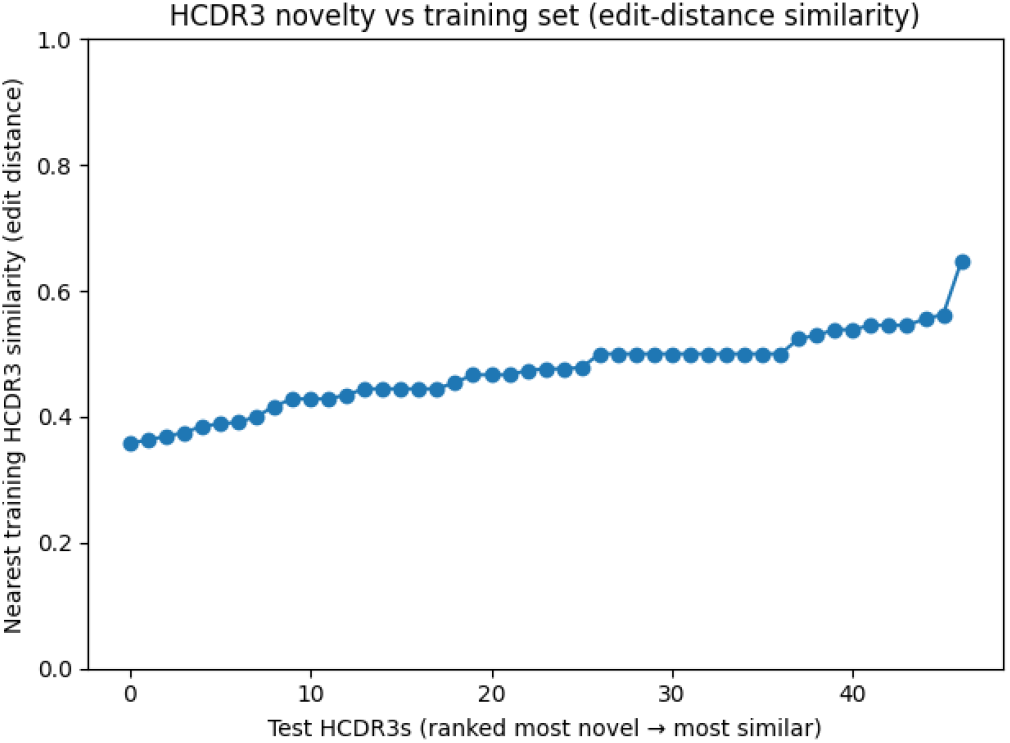
HCDR3 nearest-neighbour similarity to the training and validation set. For each held-out test HCDR3 sequence, the maximum similarity to any training- or validation-set HCDR3 was computed using Levenshtein edit distance and normalized by the maximum HCDR3 length 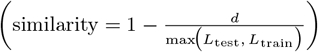 HCDR3s are ranked from most novel (left) to most similar (right).

**Figure S2:**
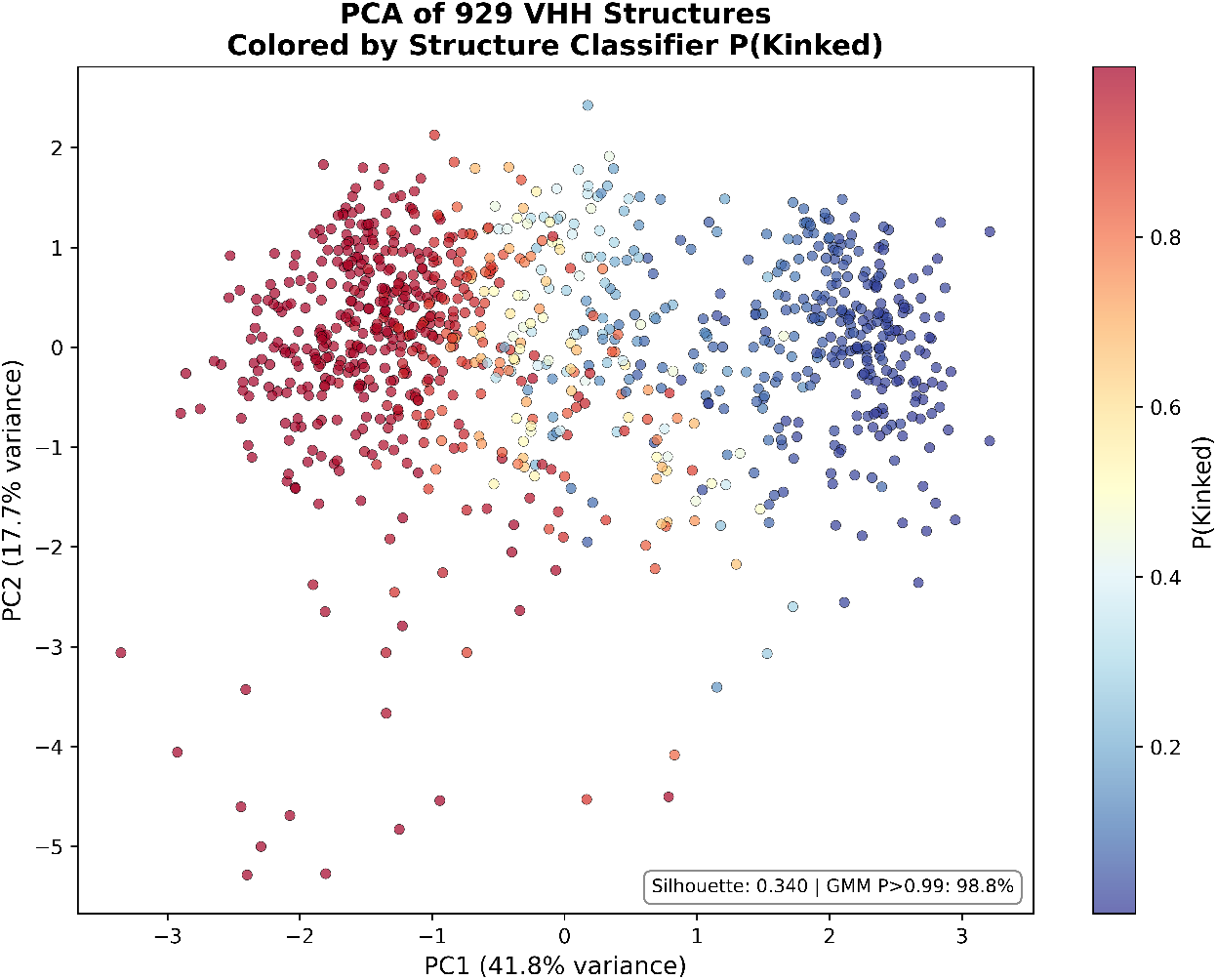
Principal Component Analysis (PCA) of 929 nanobody structures based on six structural features: N- and C-terminal CDR3 backbone angles (*α*_*N*_, *τ*_*N*_, *α*_*C*_, *τ*_*C*_), contact density (CDR3-FR2 contacts normalized by CDR3 length), and relative solvent accessibility of key FR2 positions (AHo 44 and 54). Points are coloured by the predicted probability of kinked conformation P(Kinked) from a logistic regression classifier trained on 100 expert-labelled structures from the training set. The two principal components explain 59.5% of the total variance (PC1: 41.8%, PC2: 17.7%). Separation between kinked (red, high P(Kinked)) and extended (blue, low P(Kinked)) conformations is observed, with a small population of intermediate structures in the transition zone. The moderate silhouette score (0.340) reflects the continuum between kinked and extended conformations, where sharp boundaries are not expected. However, Gaussian Mixture Model (GMM) analysis demonstrates that 98.8% of structures can be assigned to one of two clusters with confidence >0.99, indicating that the six-dimensional feature space provides discriminative power despite geometric overlap in the 2D projection.

**Figure S3:**
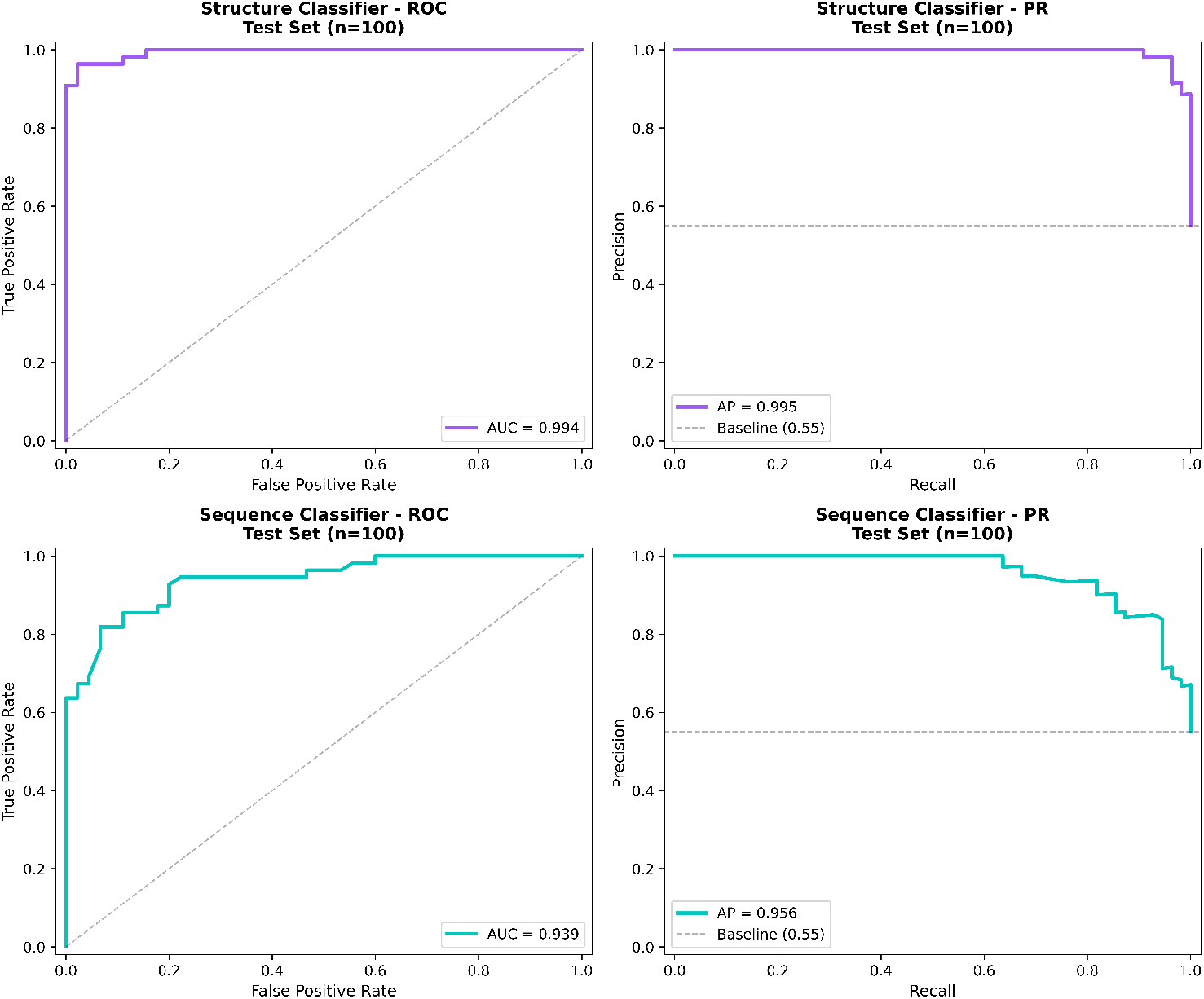
Performance evaluation of NbFrame classifiers on 100 held-out nanobody structures (PDB IDs starting with “9”) with manual labels as ground truth. Top row: Structure-based classifier using six geometric features (N- and C-terminal CDR3 backbone angles, CDR3-FR2 contact density, and FR2 solvent accessibility at key positions). Bottom row: Sequence-based classifier using 20 framework hallmark features from 10 positions. Left column: Receiver Operating Characteristic (ROC) curves showing true positive rate versus false positive rate; dashed diagonal indicates random classification. Right column: Precision-Recall (PR) curves; dashed horizontal line indicates the baseline precision equal to the proportion of kinked structures in the test set (0.55). The structure classifier achieves near-perfect discrimination (ROC-AUC = 0.994, AP = 0.995), while the sequence classifier, using only amino acid sequence information, achieves strong performance (ROC-AUC = 0.939, AP = 0.956).

**Figure S4:**
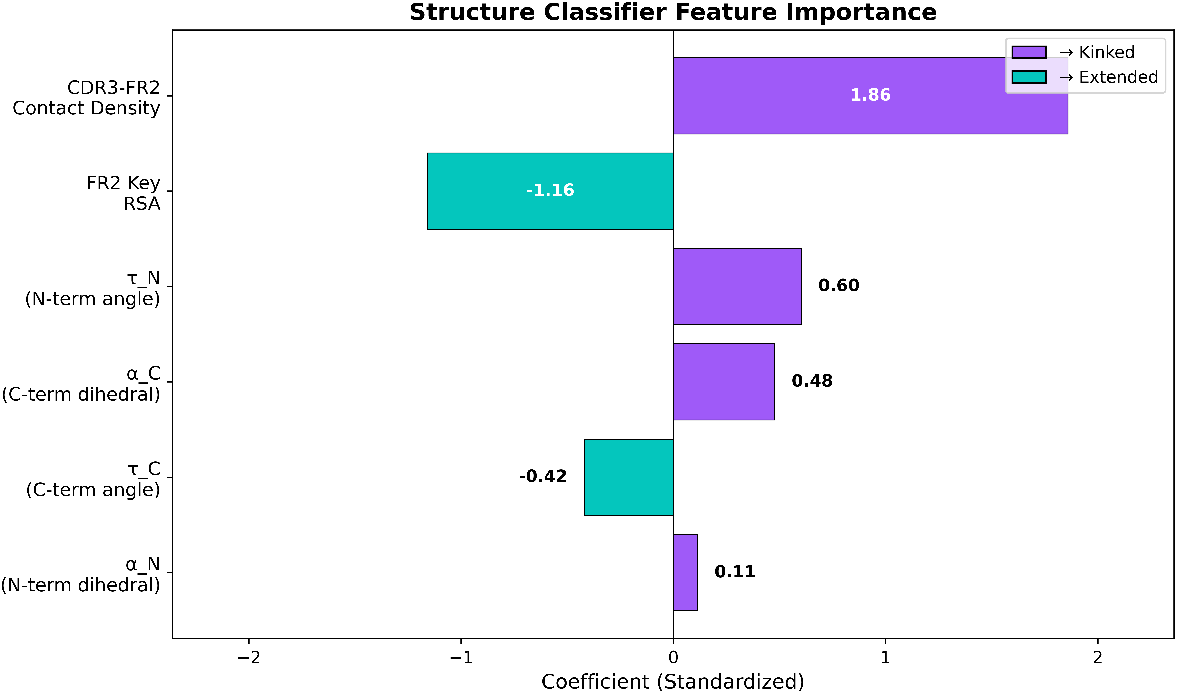
Structure classifier feature importance. Standardized logistic regression coefficients for the six structural features used in binary classification. Positive coefficients (purple) indicate features associated with kinked conformations, while negative coefficients (cyan) indicate features associated with extended conformations. CDR3-FR2 contact density (contacts normalized by CDR3 length at FR2 positions 44-55) is the strongest predictor of kinked conformation (coefficient: +1.86), followed by FR2 key position relative solvent accessibility (RSA at positions 44 and 54, coefficient: -1.16), where lower RSA indicates buried FR2 residues characteristic of kinked structures. The four backbone geometry features (*τ*_*N*_, *α*_*C*_, *τ*_*C*_, *α*_*N*_) contribute less to classification (coefficients: +0.60, +0.48, -0.42, +0.11), indicating that CDR3-FR2 packing interactions are more discriminative than CDR3 geometry alone. The trained classifier achieves ROC-AUC of 0.994 and 94% accuracy on the test set (n=100).

**Figure S5:**
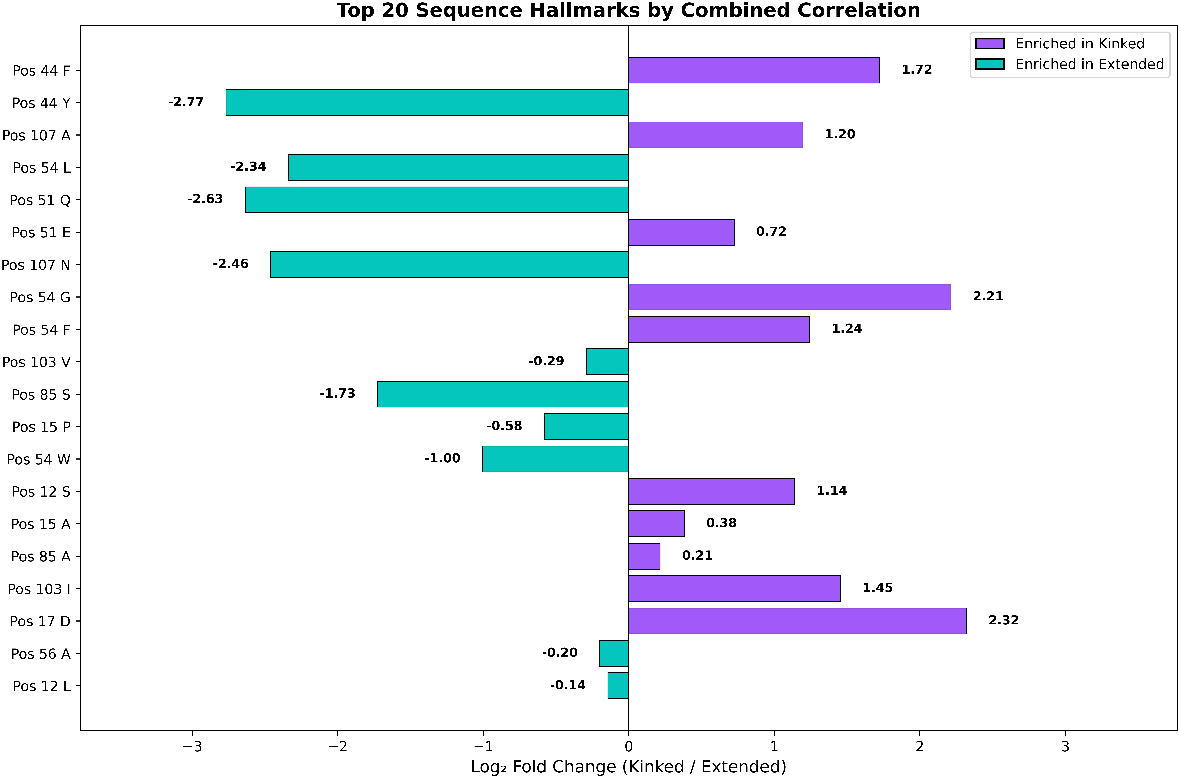
Top 20 sequence hallmarks for kinked/extended classification (AHo numbering). Log_2_ fold change (Kinked/Extended) for position-specific amino acid features selected by combined correlation with both binary labels and structure classifier predictions. Positive values (purple) indicate amino acids enriched in kinked conformations, while negative values (cyan) indicate enrichment in extended conformations. Hallmarks are drawn from framework regions and the HCDR3 N-terminal anchor position, concentrated at the CDR3-framework interface: FR2 positions 44, 51, 54, 56 (9 features), FR3 positions 85 and 103 (4 features), FR1 positions 12, 15, 17 (5 features), and position 107 at the HCDR3 N-terminal stem (2 features). Position 107, immediately C-terminal to the conserved Cys106, anchors the HCDR3 loop to the framework; unlike hypervariable CDR positions, it shows limited but functionally meaningful sequence variation associated with loop conformation. The strongest discriminative features are at FR2 position 44, where phenylalanine (F, log_2_FC: +1.72) strongly predicts kinked conformation and tyrosine (Y, log_2_FC: *−*2.77) predicts extended conformation. Other key hallmarks include position 107 alanine (+1.20, kinked), position 54 glycine (+2.21, kinked), and position 51 glutamine (*−*2.63, extended). The sequence classifier trained on these 20 hallmarks achieves ROC-AUC of 0.939 and 86% accuracy on the test set (*n* = 100), with strong correlation to the structure classifier (Pearson r=0.812).

**Figure S6:**
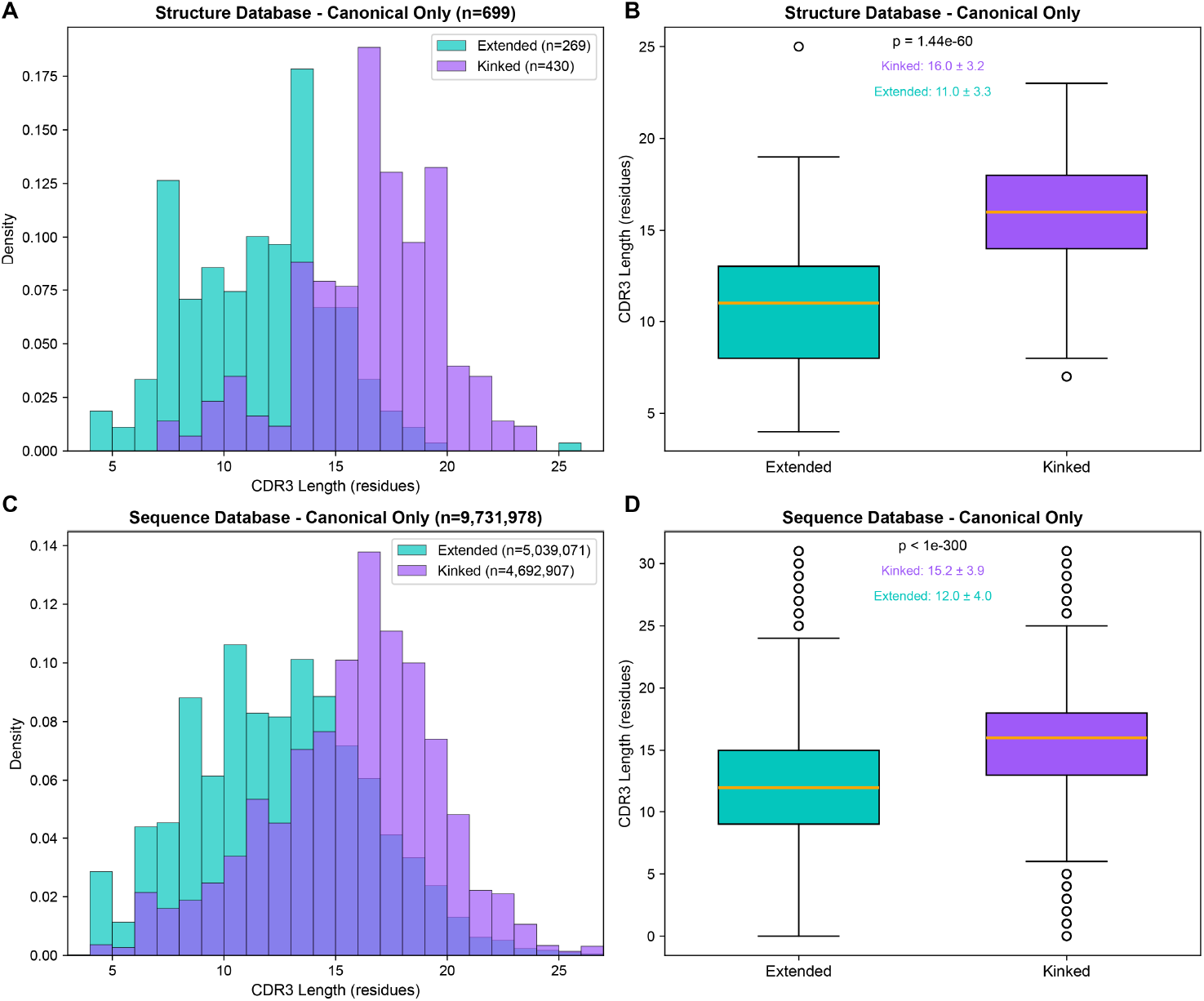
Relationship between CDR3 length and predicted conformation in nanobodies with canonical disulphide bonds. Analysis was restricted to sequences containing exactly two cysteine residues to exclude nanobodies with non-canonical disulphide bonds, which tend to have longer CDR3 loops and are predominantly kinked. Only confident predictions were included, excluding sequences in the classifier uncertainty zones. **(A-B):** Structure database (*n* = 699). Predicted kinked conformations (purple, *n* = 430) have significantly longer CDR3 loops (mean 16.0 ± 3.2 residues) compared to predicted extended conformations (teal, *n* = 269; mean 11.0 ±3.3 residues; Mann-Whitney U test, p = 1.44 *×* 10^*−*60^). **(C-D):** Sequence database (*n* = 9, 731, 978). The same trend is observed at scale, with predicted kinked sequences (*n* = 4, 692, 907) having longer CDR3 loops (mean 15.2 ± 3.9 residues) than predicted extended sequences (*n* = 5, 039, 071; mean 12.0 ± 4.0 residues; p < 10^*−*300^). Default classifiers thresholds were used: structure classifier (extended: P<0.25, kinked: P>0.55); sequence classifier (extended: P<0.40, kinked: P>0.70).

**Figure S7:**
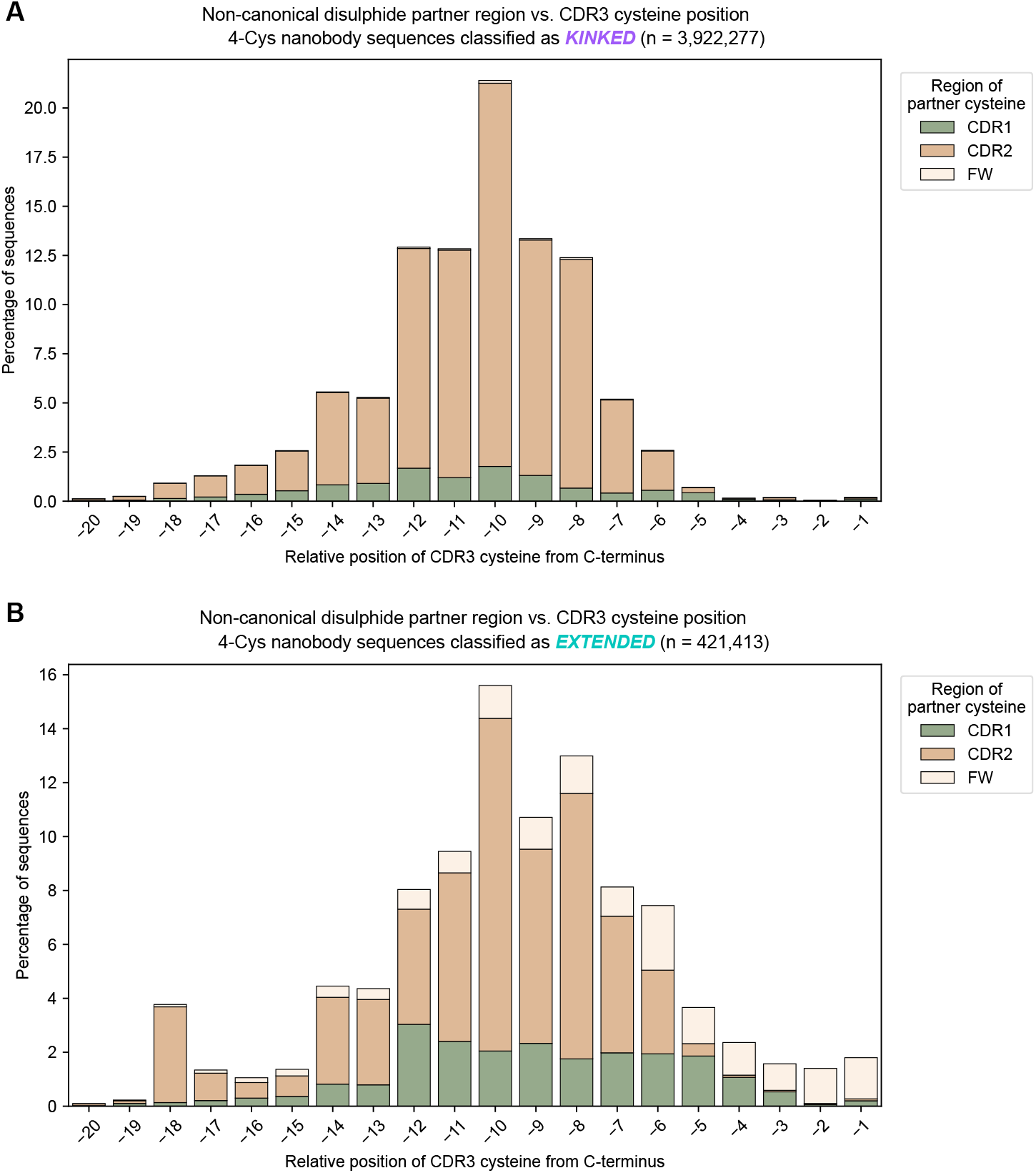
Structural region of the putative non-canonical disulphide partner cysteine as a function of CDR3 cysteine position. Analysis was restricted to sequences containing exactly four cysteine residues: the canonical disulphide pair (AHo positions 23 and 106), exactly one additional cysteine within CDR3 (AHo positions 108–138), and one partner cysteine elsewhere in the sequence, presumably forming a non-canonical disulphide bond with the CDR3 cysteine. The x-axis indicates the relative position of the CDR3 cysteine from the C-terminus of CDR3 (AHo position 138 = *−*1), counting only non-gap residues. Stacked bars are coloured by the structural region of the partner cysteine: CDR1 (AHo positions 27–42, green), CDR2 (AHo positions 57–69, tan), or framework (all other non-CDR3 positions, cream). **(A)** Sequences predicted as kinked (*n* = 3, 922, 277). The partner cysteine is overwhelmingly located in CDR2, with a minor contribution from CDR1 and nearly no framework partners, consistent with the well-characterised CDR2–CDR3 non-canonical disulphide bond that stabilises the kinked CDR3 conformation. The CDR3 cysteine position shows a pronounced peak at approximately − 10 from the C-terminus. **(B)** Sequences predicted as extended (*n* = 421, 413). CDR2 remains the predominant partner region but with substantially greater contributions from both CDR1 and framework compared to kinked sequences. The CDR3 cysteine positions are more broadly distributed across the CDR3 loop, suggesting more diverse non-canonical disulphide bond geometries in the extended conformation. Default sequence classifier thresholds were used (extended: P<0.40, kinked: P>0.70).

**Figure S8:**
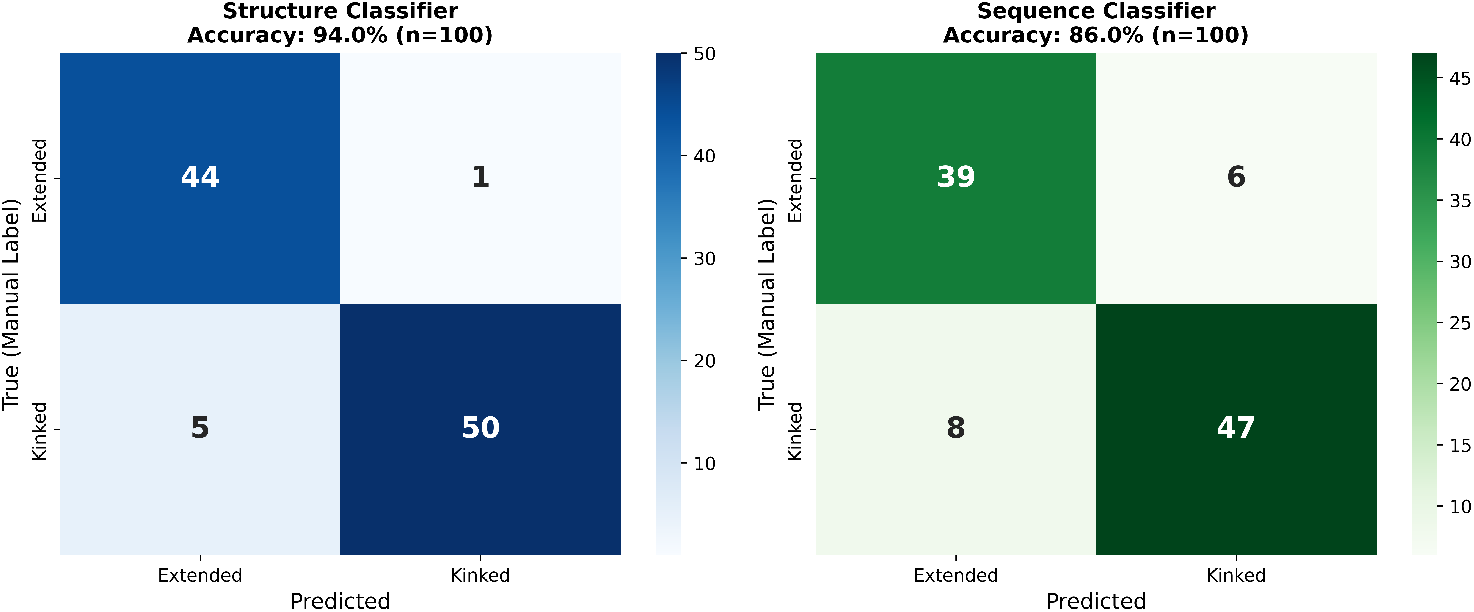
Test set confusion matrices for both classifiers. Confusion matrices showing classification performance on 100 held-out test structures (PDB IDs starting with ‘9’). **Left:** Structure classifier achieves 94% accuracy with high precision for both classes. True negatives (Extended correctly classified): 44/45 (97.8%). True positives (Kinked correctly classified): 50/55 (90.9%). Only 6 total misclassifications: 1 false positive and 5 false negatives, all occurring in structures with intermediate predicted probabilities (0.3 < P < 0.7). **Right:** Sequence classifier achieves 86% accuracy with balanced performance. True negatives: 39/45 (86.7%). True positives: 47/55 (85.5%). Total misclassifications: 14 (6 false positives, 8 false negatives). The sequence classifier’s error distribution is more symmetric, reflecting the greater challenge of predicting 3D conformation from sequence information alone. Both classifiers show balanced precision and recall, with no systematic bias toward either class.

**Figure S9:**
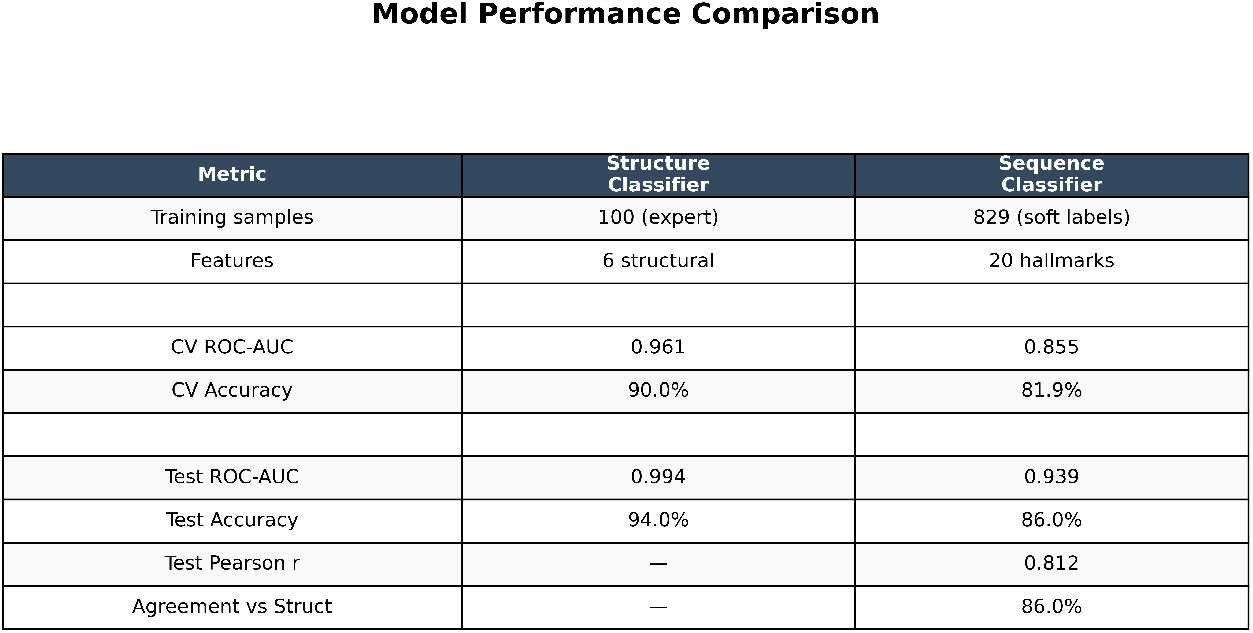
NbFrame classifier performance comparison. Summary of key performance metrics for the NbFrame structure and sequence classifiers evaluated on the same 100-structure test set (PDB IDs starting with ‘9’). The structure classifier was trained on 100 manually labelled structures using 6 structural features (CDR3 backbone angles, CDR3-FR2 contact density, and FR2 key position RSA), achieving cross-validation ROC-AUC of 0.961 (90% accuracy) and test set ROC-AUC of 0.994 (94% accuracy). The sequence classifier was trained on all 829 training structures with soft probability labels from the structure classifier, using 20 framework sequence hallmarks selected by combined correlation. It achieves cross-validation ROC-AUC of 0.855 (81.9% accuracy) and test set ROC-AUC of 0.939 (86% accuracy). The sequence classifier shows strong correlation with the structure classifier (Pearson r=0.812) and 86% classification agreement on the test set. Both classifiers demonstrate excellent generalization, with test performance meeting or exceeding cross-validation performance. The structure classifier’s superior performance reflects direct access to 3D structural information, while the sequence classifier’s strong performance demonstrates that CDR3 conformation can be predicted from framework sequence hallmarks alone with high confidence.

**Figure S10:**
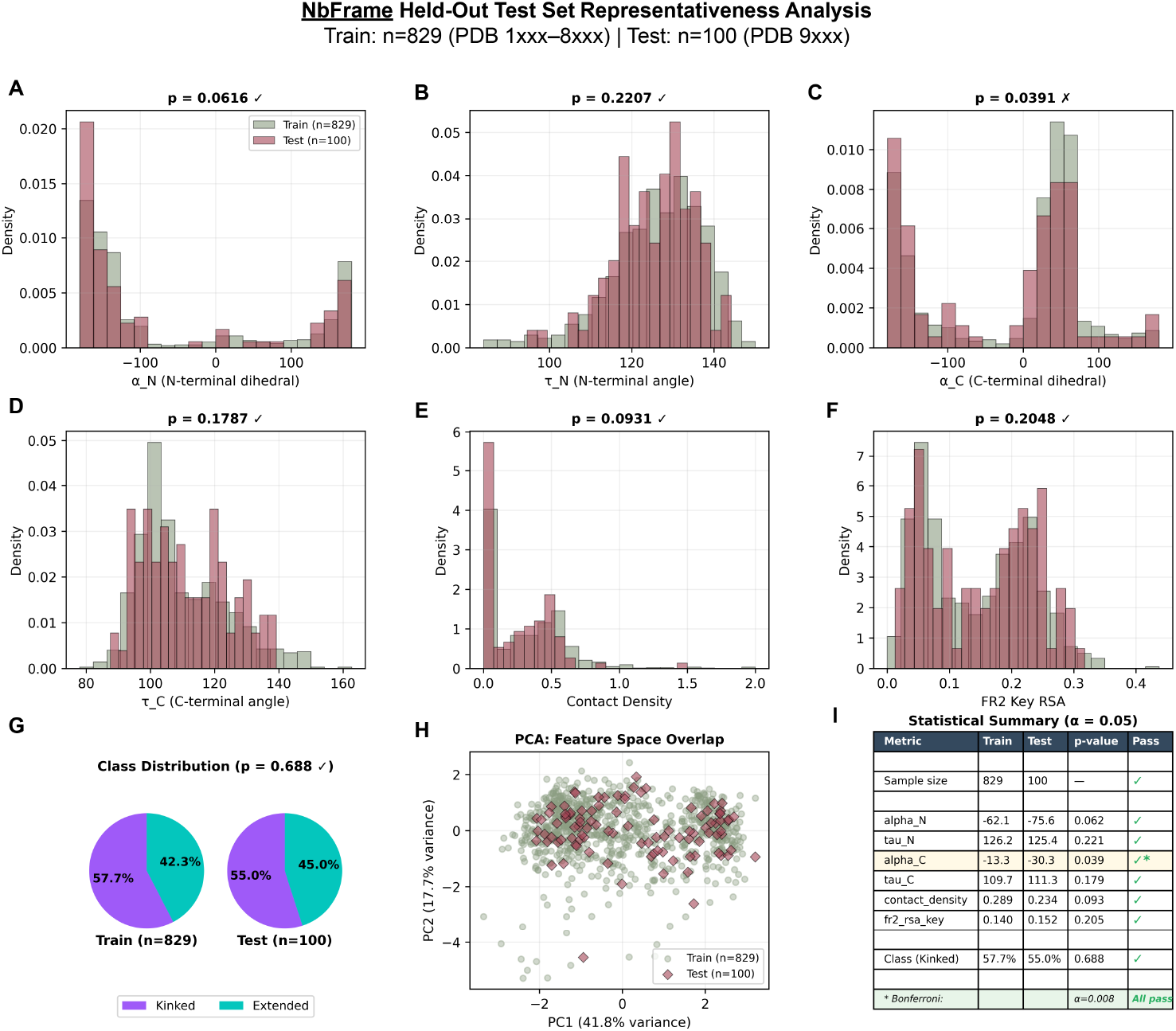
NbFrame Held-out test set representativeness analysis. Statistical comparison between NbFrame training (*n* = 829, PDB 1xxx–8xxx, green) and test (*n* = 100, PDB 9xxx, red) sets to verify unbiased temporal split. **(A–F)** Distribution comparisons for all six structural features using overlapping histograms with Mann-Whitney U test p-values. Five of six features show no significant difference (p>0.05): alpha_N (p=0.062), tau_N (p=0.221), tau_C (p=0.179), contact_density (p=0.093), and fr2_rsa_key (p=0.205). The alpha_C feature shows marginal significance (p=0.039) at the uncorrected threshold; however, when testing multiple features (*n* = 6), the Bonferroni-corrected significance threshold is *α*=0.05/6=0.008, and all features including alpha_C pass this more stringent criterion. This correction accounts for the increased risk of false positives when performing multiple statistical tests simultaneously. Additionally, alpha_C has minimal impact on classification (coefficient rank 4/6, standardized coefficient +0.477) with the model primarily driven by contact_density (+1.859) and fr2_rsa_key (*−*1.158). **(G)** Class distribution comparison via side-by-side pie charts shows similar proportions of Kinked structures in training (57.7%, 478/829) and test (55.0%, 55/100) sets (chi-squared test p=0.688), confirming balanced class representation. **(H)** Principal component analysis of the six-dimensional feature space demonstrates substantial overlap between training (green circles) and test (red diamonds) distributions. PC1 and PC2 explain 41.8% and 10.7% of variance respectively, with test structures distributed throughout the training feature space rather than clustered separately, indicating the test set samples from the same underlying distribution. **(I)** Statistical summary table showing all features with green checkmarks, including alpha_C marked with an asterisk (*) to denote marginal p-value at the uncorrected threshold. The bottom row notes that with Bonferroni correction (*α*=0.008), all features pass the representativeness criterion. Overall, the held-out test set is representative of the training set across structural features, class proportions, and feature space geometry, validating the temporal split strategy for unbiased model evaluation and supporting generalization of classifier performance to unseen structures.

**Figure S11:**
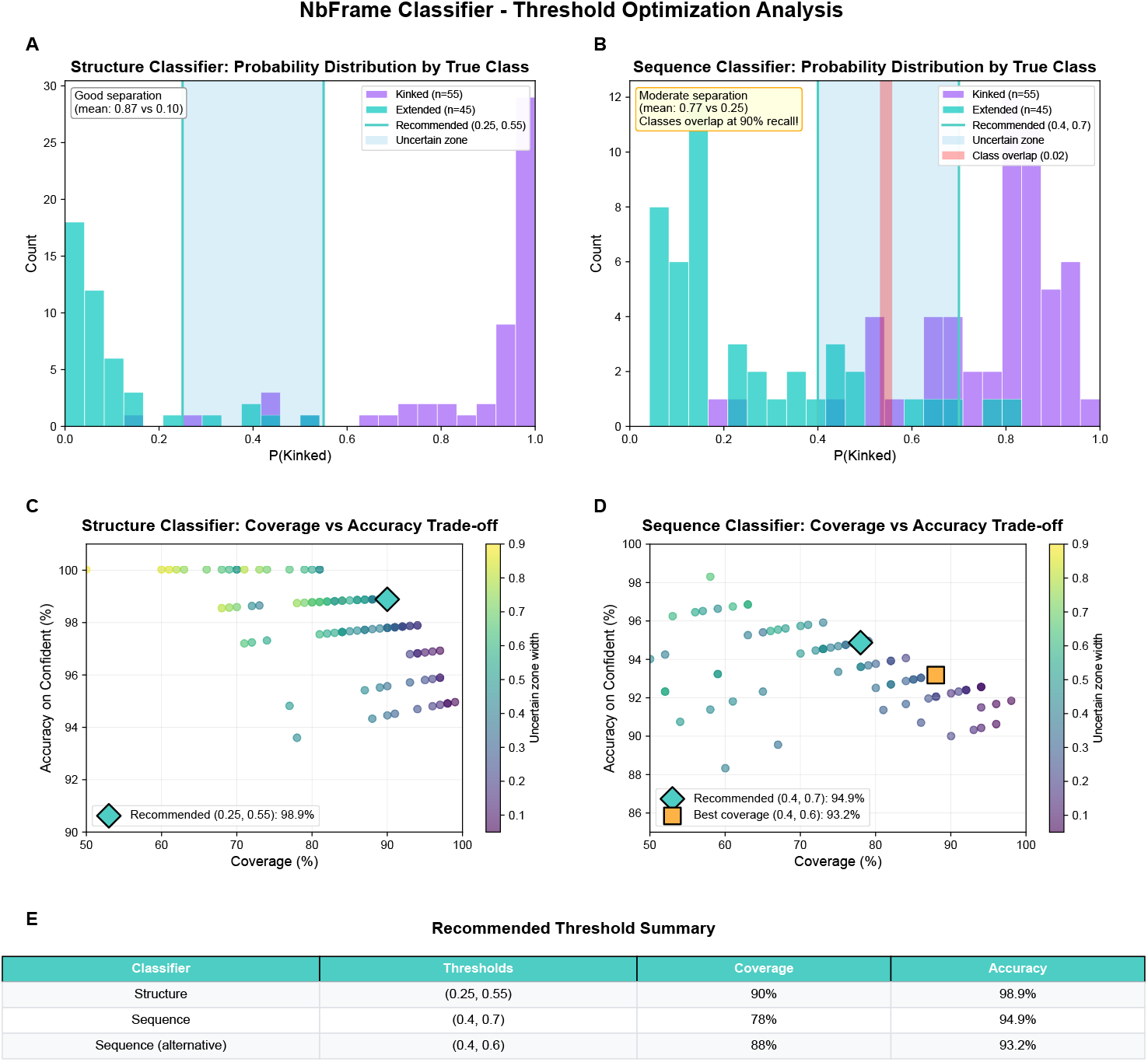
NbFrame classifier threshold optimization analysis. Systematic evaluation of classification thresholds for the structure-based and sequence-based classifiers on the held-out test set (*n* = 100 manually labelled structures: 55 Kinked, 45 Extended). **(A)** Structure classifier probability distribution by true class shows excellent separation between Kinked (purple, mean P(Kinked)=0.87) and Extended (teal, mean P(Kinked)=0.10) structures, with a class mean gap of 0.77. Recommended thresholds (0.25, 0.55) define the uncertain zone (blue shading) where predictions are withheld due to ambiguity. **(B)** Sequence classifier probability distribution shows moderate separation between classes (Kinked mean=0.77, Extended mean=0.25), with notable class overlap (red shading) at the 90% recall threshold, reflecting the fundamental limitation of sequence-only prediction compared to structure-based classification. Recommended thresholds (0.4, 0.7) balance accuracy and coverage. **(C)** Structure classifier coverage versus accuracy trade-off across all threshold combinations tested via grid search. Points are colored by uncertain zone width. The recommended thresholds (0.25, 0.55; diamond marker) achieve 90% coverage with 98.9% accuracy on confident predictions, representing an optimal balance between classification certainty and sample utilization. **(D)** Sequence classifier coverage versus accuracy trade-off. The recommended thresholds (0.4, 0.7; diamond marker) achieve 78% coverage with 94.9% accuracy. An alternative configuration (0.4, 0.6; square marker) prioritizes coverage (88%) with slightly reduced accuracy (93.2%) for applications where maximum sample utilization is preferred. **(E)** Summary Table: Recommended (default) threshold configurations for both classifiers. The structure classifier achieves near-perfect accuracy (98.9%) while maintaining high coverage (90%). The sequence classifier offers two operating points: the default (0.4, 0.7) optimizes accuracy at 94.9% with 78% coverage, while the alternative (0.4, 0.6) maximizes coverage at 88% with 93.2% accuracy.

**Table S1:** RMSD comparisons for NbForge versus baselines on the NbForge test set (*n* = 47). RMSDs are backbone-atom RMSD (Å) after framework superimposition. Δ values are mean paired differences (NbForge minus different baseline models; negative values indicate improvement). P-values are from two-sided paired Wilcoxon signed-rank tests, paired *t*-tests, and paired sign-flip permutation tests on the median paired difference (20,000 permutations).

| Region | NbForge mean RMSD | $\Delta$ (NbForge – model RMSD) | | | p-value | | | | | | | | |
| --- | --- | --- | --- | --- | --- | --- | --- | --- | --- | --- | --- | --- | --- |
| | | NBB2 | AF3 | Boltz1 | Wilcoxon | | | $t$-test | | | Permutation (median) | | |
|  |  |  |  |  | NBB2 | AF3 | Boltz1 | NBB2 | AF3 | Boltz1 | NBB2 | AF3 | Boltz1 |
| All backbone | 1.57 | -0.21 | 0.10 | 0.01 | 1.4e-04 | 0.27 | 0.95 | 1.5e-04 | 0.25 | 0.88 | 1.5e-03 | 0.65 | 0.65 |
| Framework | 0.79 | -0.02 | 0.09 | 0.10 | 0.61 | <1e-04 | <1e-04 | 0.33 | <1e-04 | <1e-04 | 0.76 | <1e-04 | <1e-04 |
| HCDR1 | 1.65 | -0.22 | 0.09 | -0.01 | 0.026 | 0.33 | 0.77 | 0.028 | 0.58 | 0.97 | 0.014 | 0.15 | 0.44 |
| HCDR2 | 1.36 | 0.01 | 0.30 | 0.15 | 0.92 | <1e-04 | 0.048 | 0.83 | 1.1e-04 | 0.056 | 0.77 | 1.3e-03 | 0.042 |
| HCDR3 | 3.40 | -0.64 | 0.15 | 0.01 | 2.7e-04 | 0.49 | 0.90 | 2.7e-04 | 0.56 | 0.97 | 6.0e-04 | 0.88 | 0.77 |

**Table S2:** RMSD comparisons for AF3 versus baselines on the test set (*n* = 47). RMSDs are backbone-atom RMSD (Å) after framework superimposition. Δ values are mean paired differences (AF3 minus different baseline models; negative values indicate improvement). P-values are from two-sided paired Wilcoxon signed-rank tests, paired *t*-tests, and paired sign-flip permutation tests on the median paired difference (20,000 permutations).

| Region | AF3 mean RMSD | $\Delta$ (AF3 – model RMSD) | | | p-value | | | | | | | | |
| --- | --- | --- | --- | --- | --- | --- | --- | --- | --- | --- | --- | --- | --- |
| | | NbForge | NBB2 | Boltz1 | Wilcoxon | | | $t$-test | | | Permutation (median) | | |
|  |  |  |  |  | NbForge | NBB2 | Boltz1 | NbForge | NBB2 | Boltz1 | NbForge | NBB2 | Boltz1 |
| All backbone | 1.47 | -0.10 | -0.30 | -0.08 | 0.27 | 0.0044 | 0.094 | 0.25 | 0.0023 | 0.25 | 0.65 | 0.014 | 0.19 |
| Framework | 0.69 | -0.09 | -0.11 | 0.01 | <1e-04 | <1e-04 | 0.47 | <1e-04 | <1e-04 | 0.58 | <1e-04 | <1e-04 | 0.44 |
| HCDR1 | 1.57 | -0.09 | -0.30 | -0.09 | 0.33 | 0.024 | 0.14 | 0.58 | 0.056 | 0.48 | 0.15 | 0.024 | 0.27 |
| HCDR2 | 1.06 | -0.30 | -0.29 | -0.16 | <1e-04 | <1e-04 | 0.14 | <1e-04 | <1e-04 | 0.026 | 0.0019 | 0.0035 | 0.76 |
| HCDR3 | 3.25 | -0.15 | -0.78 | -0.14 | 0.49 | 0.017 | 0.21 | 0.56 | 0.0076 | 0.53 | 0.88 | 0.066 | 0.44 |

**Table S3:**
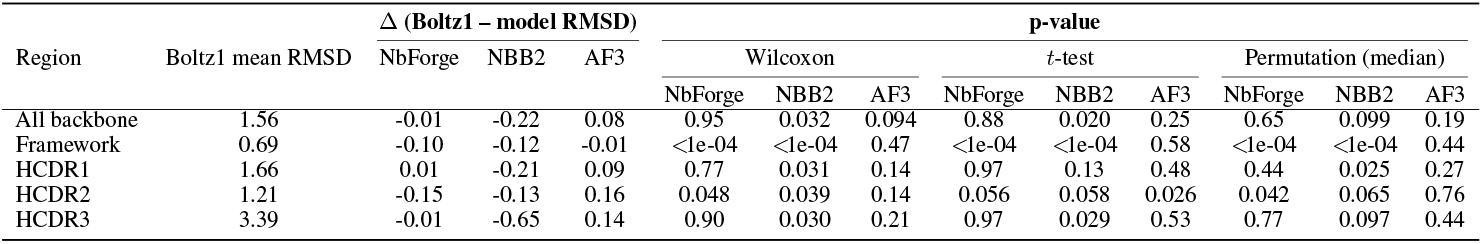
RMSD comparisons for Boltz1 versus baselines on the test set (*n* = 47). RMSDs are backbone-atom RMSD (Å) after framework superimposition. Δ values are mean paired differences (Boltz1 minus different baseline models; negative values indicate improvement). P-values are from two-sided paired Wilcoxon signed-rank tests, paired *t*-tests, and paired sign-flip permutation tests on the median paired difference (20,000 permutations).

| Region | Boltz1 mean RMSD | $\Delta$ (Boltz1 – model RMSD) | | | p-value | | | | | | | | |
| --- | --- | --- | --- | --- | --- | --- | --- | --- | --- | --- | --- | --- | --- |
| | | NbForge | NBB2 | AF3 | Wilcoxon | | | $t$-test | | | Permutation (median) | | |
|  |  |  |  |  | NbForge | NBB2 | AF3 | NbForge | NBB2 | AF3 | NbForge | NBB2 | AF3 |
| All backbone | 1.56 | -0.01 | -0.22 | 0.08 | 0.95 | 0.032 | 0.094 | 0.88 | 0.020 | 0.25 | 0.65 | 0.099 | 0.19 |
| Framework | 0.69 | -0.10 | -0.12 | -0.01 | <1e-04 | <1e-04 | 0.47 | <1e-04 | <1e-04 | 0.58 | <1e-04 | <1e-04 | 0.44 |
| HCDR1 | 1.66 | 0.01 | -0.21 | 0.09 | 0.77 | 0.031 | 0.14 | 0.97 | 0.13 | 0.48 | 0.44 | 0.025 | 0.27 |
| HCDR2 | 1.21 | -0.15 | -0.13 | 0.16 | 0.048 | 0.039 | 0.14 | 0.056 | 0.058 | 0.026 | 0.042 | 0.065 | 0.76 |
| HCDR3 | 3.39 | -0.01 | -0.65 | 0.14 | 0.90 | 0.030 | 0.21 | 0.97 | 0.029 | 0.53 | 0.77 | 0.097 | 0.44 |

**Table S4:** RMSD comparisons for NBB2 versus baselines on the test set (*n* = 47). RMSDs are backbone-atom RMSD (Å) after framework superimposition. Δ values are mean paired differences (NBB2 minus different baseline models; negative values indicate improvement). P-values are from two-sided paired Wilcoxon signed-rank tests, paired *t*-tests, and paired sign-flip permutation tests on the median paired difference (20,000 permutations).

| Region | NBB2 mean RMSD | $\Delta$ (NBB2 – model RMSD) | | | p-value | | | | | | | | |
| --- | --- | --- | --- | --- | --- | --- | --- | --- | --- | --- | --- | --- | --- |
| | | NbForge | AF3 | Boltz1 | Wilcoxon | | | $t$-test | | | Permutation (median) | | |
|  |  |  |  |  | NbForge | AF3 | Boltz1 | NbForge | AF3 | Boltz1 | NbForge | AF3 | Boltz1 |
| All backbone | 1.78 | 0.21 | 0.30 | 0.22 | 1.4e-04 | 0.0044 | 0.032 | 1.5e-04 | 0.0023 | 0.020 | 1.5e-03 | 0.014 | 0.097 |
| Framework | 0.81 | 0.02 | 0.11 | 0.12 | 0.61 | <1e-04 | <1e-04 | 0.33 | <1e-04 | <1e-04 | 0.76 | <1e-04 | <1e-04 |
| HCDR1 | 1.87 | 0.22 | 0.30 | 0.21 | 0.026 | 0.024 | 0.031 | 0.028 | 0.056 | 0.13 | 0.014 | 0.024 | 0.026 |
| HCDR2 | 1.35 | -0.01 | 0.29 | 0.13 | 0.92 | <1e-04 | 0.039 | 0.83 | <1e-04 | 0.058 | 0.77 | 0.0035 | 0.064 |
| HCDR3 | 4.03 | 0.64 | 0.78 | 0.65 | 2.7e-04 | 0.017 | 0.030 | 2.7e-04 | 0.0076 | 0.029 | 6.0e-04 | 0.066 | 0.10 |

**Table S5:** Coordinate accuracy and feature recovery on the NbForge test set. Mean Backbone (CA, N, CO, O) RMSDs are reported as mean ± standard error (SE) in Å (*n* = 47) after framework superimposition. Kinked/extended (K/E) blueprint recovery is computed over all test structures (see Methods), while non-canonical disulphide bond (NCDB) recovery is computed over the subset containing a NCDB (*n* = 15), see Methods.

| Model | All | FR | HCDR1 | HCDR2 | HCDR3 | K/E<br>rec. (%) | NCDB<br>rec. (%) |
| --- | --- | --- | --- | --- | --- | --- | --- |
| AF3 | $1.47 \pm 0.10$ | $0.69 \pm 0.04$ | $1.57 \pm 0.15$ | $1.06 \pm 0.09$ | $3.25 \pm 0.30$ | 93.6 | 93.3 |
| Boltz1 | $1.56 \pm 0.10$ | $0.69 \pm 0.04$ | $1.66 \pm 0.16$ | $1.21 \pm 0.13$ | $3.39 \pm 0.30$ | 89.4 | 73.0 |
| NBB2 | $1.78 \pm 0.08$ | $0.81 \pm 0.04$ | $1.87 \pm 0.12$ | $1.35 \pm 0.12$ | $4.03 \pm 0.25$ | 80.9 | 26.7 |
| NbForge | $1.57 \pm 0.07$ | $0.79 \pm 0.03$ | $1.65 \pm 0.11$ | $1.36 \pm 0.11$ | $3.40 \pm 0.24$ | 91.5 | 86.7 |

**Table S6:** Mean Backbone coordinate accuracy (RMSD, Å) stratified by presence of a non-canonical disulphide bond (NCDB *n* = 15) or by its absence (no NCDB *n* = 32). RMSDs are calculated for all backbone heavy atoms following framework superimposition and reported as median with bootstrap 95% confidence intervals.

| NCDB | Region | AF3 | Boltz1 | NBB2 | NbForge |
| --- | --- | --- | --- | --- | --- |
| no NCDB | FR | $0.71 \pm 0.05$ | $0.72 \pm 0.05$ | $0.83 \pm 0.05$ | $0.82 \pm 0.05$ |
| | HCDR1 | $1.49 \pm 0.19$ | $1.48 \pm 0.18$ | $1.83 \pm 0.16$ | $1.64 \pm 0.15$ |
| | HCDR2 | $0.98 \pm 0.11$ | $1.09 \pm 0.13$ | $1.22 \pm 0.12$ | $1.31 \pm 0.13$ |
| | HCDR3 | $3.15 \pm 0.41$ | $3.30 \pm 0.40$ | $4.11 \pm 0.33$ | $3.47 \pm 0.33$ |
| NCDB | FR | $0.65 \pm 0.04$ | $0.63 \pm 0.05$ | $0.77 \pm 0.04$ | $0.73 \pm 0.04$ |
| | HCDR1 | $1.60 \pm 0.27$ | $1.90 \pm 0.28$ | $1.94 \pm 0.19$ | $1.63 \pm 0.16$ |
| | HCDR2 | $1.24 \pm 0.16$ | $1.52 \pm 0.31$ | $1.65 \pm 0.24$ | $1.50 \pm 0.22$ |
| | HCDR3 | $3.39 \pm 0.45$ | $3.50 \pm 0.45$ | $3.92 \pm 0.39$ | $3.23 \pm 0.27$ |

**Table S7:** Coordinate accuracy and feature recovery on the NbForge test set. Mean Backbone (CA, N, CO, O) RMSDs are reported as mean ± standard error (SE) in Å (*n* = 47) after framework superimposition. Kinked/extended (K/E) blueprint recovery is computed over all test structures (see Methods), while non-canonical disulphide bond (NCDB) recovery is computed over the subset containing a NCDB (*n* = 15), see Methods. NbForge models without OpenMM relaxation are used in these calculations.

| Model | All | FR | HCDR1 | HCDR2 | HCDR3 | K/E<br>rec. (%) | NCDB<br>rec. (%) |
| --- | --- | --- | --- | --- | --- | --- | --- |
| AF3 | $1.47 \pm 0.10$ | $0.69 \pm 0.04$ | $1.57 \pm 0.15$ | $1.06 \pm 0.09$ | $3.25 \pm 0.30$ | 93.6 | 93.3 |
| Boltz1 | $1.56 \pm 0.10$ | $0.69 \pm 0.04$ | $1.66 \pm 0.16$ | $1.21 \pm 0.13$ | $3.39 \pm 0.30$ | 89.4 | 73.0 |
| NBB2 | $1.78 \pm 0.08$ | $0.81 \pm 0.04$ | $1.87 \pm 0.12$ | $1.35 \pm 0.12$ | $4.03 \pm 0.25$ | 80.9 | 26.7 |
| NbForge | $1.57 \pm 0.07$ | $0.79 \pm 0.03$ | $1.65 \pm 0.11$ | $1.35 \pm 0.11$ | $3.38 \pm 0.23$ | 89.4 | 80.0 |

**Table S8:** Coordinate accuracy and feature recovery on the NbForge test set. Backbone (CA, N, CO, O) RMSDs are reported as median with bootstrap 95% confidence intervals (CI) in Å (*n* = 47) after framework superimposition. Confidence intervals were estimated using non-parametric bootstrap resampling with 10,000 resamples. Kinked/extended (K/E) correctness is computed over all test structures, while non-canonical disulphide bond (NCDB) recovery is computed over the subset containing a non-canonical disulphide bond (*n* = 15), see Methods. NbForge models without OpenMM relaxation are used in these calculations.

| Metric | AF3 | Boltz1 | NBB2 | NbForge |
| --- | --- | --- | --- | --- |
| All | 1.39 [1.05, 1.53] | 1.52 [1.23, 1.76] | 1.62 [1.51, 1.91] | 1.47 [1.34, 1.63] |
| FR | 0.70 [0.57, 0.74] | 0.65 [0.58, 0.73] | 0.79 [0.67, 0.85] | 0.73 [0.69, 0.78] |
| HCDR1 | 1.19 [1.03, 1.50] | 1.28 [1.02, 1.63] | 1.72 [1.53, 2.17] | 1.57 [1.36, 1.77] |
| HCDR2 | 0.80 [0.67, 1.26] | 0.89 [0.63, 1.33] | 1.17 [0.94, 1.44] | 1.17 [1.02, 1.43] |
| HCDR3 | 2.95 [2.30, 3.66] | 3.39 [2.47, 3.74] | 3.45 [2.99, 4.53] | 2.97 [2.72, 3.63] |
| K/E recovery (%) | 93.6 | 89.4 | 80.9 | 89.4 |
| NCDB recovery (%) | 93.3 | 73.0 | 26.7 | 80.0 |

## References

[1] C. Hamers-Casterman, T. Atarhouch, S. Muyldermans, G. Robinson, C. Hammers, E. Bajyana Songa, N. Bendahman, and R. Hammers. Naturally occurring antibodies devoid of light chains. Nature, 363(6428):446–448, 1993. doi: 10.1038/363446a0.

[2] Sean Duggan. Caplacizumab: First global approval. Drugs, 78(15):1639–1642, 2018. doi: 10.1007/s40265-018-0989-0.

[3] Marie Scully, Spero R. Cataland, Flora Peyvandi, Paul Coppo, Paul Knöbl, Johanna A. Kremer Hovinga, Ara Metjian, Javier De La Rubia, Katerina Pavenski, Filip Callewaert, Debjit Biswas, Hilde De Winter, Robert K. Zeldin, and HERCULES Investigators. Caplacizumab treatment for acquired thrombotic thrombocytopenic purpura. New England Journal of Medicine, 380(4):335–346, 2019. doi: 10.1056/NEJMoa1806311.

[4] Michael Mullin, James McClory, Winston Haynes, Justin Grace, Nathan Robertson, and Gino van Heeke. Applications and challenges in designing vhh-based bispecific antibodies: leveraging machine learning solutions. mAbs, 16(1):2341443, 2024. doi: 10.1080/19420862.2024.2341443.

[5] Mohammed Al-Seragi, Yilun Chen, and Franck Duong van Hoa. Advances in nanobody multimerization and multispecificity: from in vivo assembly to in vitro production. Biochemical Society Transactions, 53(1):235–248, 2025. doi: 10.1042/BST20241419. URL https://www.ncbi.nlm.nih.gov/pmc/articles/PMC12203927/.

[6] Marcus J. Rohovie, Maya Nagasawa, and James R. Swartz. Virus-like particles: Next-generation nanoparticles for targeted therapeutic delivery. Bioengineering & Translational Medicine, 2 (1):43–57, 2017. doi: 10.1002/btm2.10049. URL https://www.ncbi.nlm.nih.gov/pmc/articles/PMC5689521/.

[7] Pol Escudé Martinez de Castilla, Vincenzo Verdi, Willemijn de Voogt, Mariona Estapé Sentí, Arnold C. Koekman, Julian Rietveld, Sven van Kempen, Qiangbing Yang, Juliette van Merris, Guido Jenster, Martin E. van Royen, Marcel H. Fens, Sander A. A. Kooijmans, Wytske M. van Weerden, Guillaume van Niel, Pieter Vader, and Raymond M. Schiffelers. Nanobody-decorated lipid nanoparticles for enhanced mrna delivery to tumors in vivo. Advanced Healthcare Materials, 14(24):2500605, 2025. doi: 10.1002/adhm.202500605. URL https://doi.org/10.1002/adhm.202500605.

[8] Pouya Safarzadeh Kozani, Abdolhossein Naseri, Seyed Mohamad Javad Mirarefin, Faeze Salem, Mojtaba Nikbakht, Sahar Evazi Bakhshi, and Pooria Safarzadeh Kozani. Nanobody-based car-t cells for cancer immunotherapy. Journal of Translational Medicine, 20(1):487, 2022. doi: 10.1186/s12967-022-03672-1. URL https://www.ncbi.nlm.nih.gov/pmc/articles/PMC9036779/.

[9] Shasha Guo and Xiaozhi Xi. Nanobody-enhanced chimeric antigen receptor t-cell therapy: overcoming barriers in solid tumors with vhh and vnar-based constructs. Biomarker Research, 13(1):41, 2025. doi: 10.1186/s40364-025-00755-5. URL https://biomarkerres.biomedcentral.com/articles/10.1186/s40364-025-00755-5.

[10] Angelo Rosace, A. Bennett, M. Oeller, M. M. Mortensen, L. Sakhnini, N. Lorenzen, C. Poulsen, and P. Sormanni. Automated optimisation of solubility and conformational stability of antibodies and proteins. Nature Communications, 14:1937, 2023. doi: 10.1038/s41467-023-37668-6. URL https://www.nature.com/articles/s41467-023-37668-6.

[11] Eva Smorodina, Oliver Crook, Johannes R. Loeffler, Monica Lisa Fernandez-Quintero, Lucas Matthias Weissenborn, Hannah L. Turner, Aleksandar Antanasijevic, Rahmad Akbar, Puneet Rawat, Khang Løe Quý, Brij Bhushan Mehta, Ole Magnus Fløgstad, Dario Segura-Peña, Nikolina Sekulić, Andrew B. Ward, Fridtjof Lund-Johansen, Jan Terje Andersen, and Victor Greiff. Structural modeling of antibody variant epitope specificity with complementary experimental and computational techniques. In Proceedings of the ICLR 2025 Workshop on Integrating Generative and Experimental Platforms for Biomolecular Design, 2025. URL https://openreview.net/forum?id=d6ExTOu00A.

[12] Koichi Yamamoto, Satoru Nagatoishi, Ryo Matsunaga, Makoto Nakakido, Daisuke Kuroda, and Kouhei Tsumoto. Affinity-stability trade-off mechanism of residue 35 in framework region 2 of vhh antibodies with β-hairpin cdr3. Protein Science, 34(4):e70095, 2025. doi: 10.1002/pro.70095. URL https://onlinelibrary.wiley.com/doi/10.1002/pro.70095.

[13] John Jumper, Richard Evans, Alexander Pritzel, Tim Green, Michael Figurnov, Olaf Ronneberger, Kathryn Tunyasuvunakool, Russ Bates, Augustin Žídek, Anna Potapenko, Alex Bridgland, Clemens Meyer, Stefan A. A. Kohl, Andrew J. Ballard, Andrew Cowie, Bernardino Romera-Paredes, Stanislav Nikolov, Ravinder Jain, Jonathan Adler, Trevor Back, Stephen Petersen, Dandelion Reiman, Elena Clancy, Michal Zielinski, Martin Steinegger, Marta Pacholska, Thomas Berghammer, Sebastian Bodenstein, David Silver, Oriol Vinyals, Andrew W. Senior, Koray Kavukcuoglu, Pushmeet Kohli, and Demis Hassabis. Highly accurate protein structure prediction with alphafold. Nature, 596(7873):583–589, 2021. doi: 10.1038/s41586-021-03819-2.

[14] Zeming Lin, Halil Akin, Zaixi Wu, Joseph Vega, Mykhailo Veder, Kathryn Tunyasuvunakool, Andrew W. Senior, Mark Korek, Tim Green, Chong Qin, Shay Artzi, Demis Hassabis, John Jumper, Stefan A. A. Kohl, Chakravarthy Pas, Frank DiMaio, David Baker, Ram Rao, Tom Sercu, and Alexis Rives. Evolutionary-scale prediction of atomic-level protein structure with a language model. Nature Methods, 20(9):1235–1242, 2023. doi: 10.1038/s41592-023-01953-3.

[15] Minkyung Baek, Frank DiMaio, Ivan Anishchenko, Justas Dauparas, Sergey Ovchinnikov, Grace R. Lee, Justin Wang, Qi Cong, Lisa N. Kinch, Robert D. Schaeffer, Cristina Millán, Hyeon Park, Charles Adams, Colin R. Glassman, Adam Derry, Alexander Finkelstein, Keith Moffat, Eduardo Fernandez, John D. Westbrook, Do Hoon Kim, Jared H. Davis, and David Baker. Accurate prediction of protein structures and interactions using a three-track neural network. Science, 376(6587):abj8754, 2022. doi: 10.1126/science.abj8754.

[16] Chen Jing, Stefan Eismann, Pranav Soni, and Ron O. Dror. Fast and differentiable protein structure prediction for machine learning. Bioinformatics, 39(Supplement_1):btac637, 2023. doi: 10.1093/bioinformatics/btac637.

[17] Srivamshi Pittala and Chris Bailey-Kellogg. Learning context-aware structural representations to predict antigen and antibody binding interfaces. Bioinformatics, 36(13):3996–4003, 2020. doi: 10.1093/bioinformatics/btaa263.

[18] Matthew I. J. Raybould, Claire Marks, Konrad Krawczyk, Charlotte M. Deane, Jakob Nowak, Andrew P. Lewis, Alexander Bujotzek, and Jiye Shi. Five computational developability guidelines for therapeutic antibody profiling. Proceedings of the National Academy of Sciences, 116 (10):4025–4030, 2019. doi: 10.1073/pnas.1810576116.

[19] Edgar Liberis, Petar Veličković, Pietro Sormanni, Michele Vendruscolo, and Pietro Liò. Parapred: Antibody paratope prediction using convolutional and recurrent neural networks. Bioinformatics, 34(17):2944–2950, 2018. doi: 10.1093/bioinformatics/bty305.

[20] Andrew Waterhouse, Michele Bertoni, Stefan Bienert, Geert Studer, Giulia Tauriello, Riccardo Gumienny, Florian T. Heer, Stefan de Beer, Christoph Rempfer, Lorenzo Bordoli, Raffaele Lepore, and Torsten Schwede. Swiss-model: homology modelling of protein structures and complexes. Nucleic Acids Research, 46(W1):W296–W303, 2018. doi: 10.1093/nar/gky427.

[21] Habib Bashour, Eva Smorodina, Matteo Pariset, Jahn Zhong, Rahmad Akbar, Maria Chernigovskaya, Khang Lê Quý, Igor Snapkow, Puneet Rawat, Konrad Krawczyk, Geir Kjetil Sandve, Jose Gutierrez-Marcos, Daniel Nakhaee-Zadeh Gutierrez, Jan Terje Andersen, and Victor Greiff. Biophysical cartography of the native and human-engineered antibody landscapes quantifies the plasticity of antibody developability. Communications Biology, 7:922, 2024. doi: 10.1038/s42003-024-06561-3.

[22] Jeffrey A. Ruffolo, Lee-Shin Chu, Sai Pooja Mahajan, and Jeffrey J. Gray. Fast, accurate antibody structure prediction from deep learning on massive set of natural antibodies. Nature Communications, 14(1):2389, 2023. doi: 10.1038/s41467-023-38063-x.

[23] Fatima N. Hitawala and Jeffrey J. Gray. What does alphafold3 learn about antibody and nanobody docking, and what remains unsolved? mAbs, 17(1):2545601, August 2025. doi: 10.1080/19420862.2025.2545601. URL https://doi.org/10.1080/19420862.2025.2545601.

[24] Floriane Eshak and Anne Goupil-Lamy. Advancements in nanobody epitope prediction: A comparative study of alphafold2multimer vs alphafold3. Journal of Chemical Information and Modeling, 65(4):1782–1797, 2025. doi: 10.1021/acs.jcim.4c01877.

[25] Julian Davies and Lutz Riechmann. Antibody vh domains as small recognition units. Nature Biotechnology, 13(5):475–479, 1995. doi: 10.1038/nbt0595-475.

[26] Daisuke Kuroda. Structural classification of CDR-H3 in single-domain VHH antibodies. In Antibody Engineering. Springer, 2022. doi: 10.1007/978-1-0716-2609-2_2.

[27] Serge Muyldermans, T. N. Baral, Virna C. Retarnozzo, P. De Baetselier, E. De Genst, J. Kinne, H. Leonhardt, Stefan Magez, V. K. Nguyen, H. Revets, U. Rothbauer, B. Stijlemans, S. Tillib, U. Wernery, L. Wyns, G. Hassanzadeh-Ghassabeh, and D. Saerens. Camelid immunoglobulins and nanobody technology. Veterinary Immunology and Immunopathology, 128(1-3):178–183, 2009. doi: 10.1016/j.vetimm.2008.10.299.

[28] Erwin De Genst, Karen Silence, Klaas Decanniere, Katja Conrath, Remy Loris, Jörg Kinne, Serge Muyldermans, and Lode Wyns. Molecular basis for the preferential cleft recognition by dromedary heavy-chain antibodies. Proceedings of the National Academy of Sciences of the United States of America, 103(12):4586–4591, 2006. doi: 10.1073/pnas.0505379103.

[29] Zohreh Bahrami Dizicheh et al. Vhh cdr-h3 conformation is determined by vh germline segment. Communications Biology, 2023. doi: 10.1038/s42003-023-05241-y.

[30] Shuo Qiu, Chang Liu, Guoqiang Li, Hong Lin, Limin Cao, Kaiqiang Wang, Xiudan Wang, and Jianxin Sui. Impact of noncanonical disulphide bond on thermal resistance and binding affinity of shark-derived single-domain antibodies. ACS Biomaterials Science & Engineering, 11(4):xxx–xxx, 2025. doi: 10.1021/acsbiomaterials.4c02215. URL https://pubs.acs.org/doi/10.1021/acsbiomaterials.4c02215.

[31] Natalia E. Ketaren, Peter C. Fridy, Vladimir Malashkevich, Tanmoy Sanyal, Marc Brillantes, Mary K. Thompson, Deena A. Oren, Jeffrey B. Bonanno, Andrej Šali, Steven C. Almo, Brian T. Chait, and Michael P. Rout. Unique mechanisms to increase structural stability and enhance antigen binding in nanobodies. Structure, 33:677–690, 2025. doi: 10.1016/j.str.2025.01.019s. URL https://doi.org/10.1016/j.str.2025.01.019.

[32] Baolong Xia, Ah-Ram Kim, Feimei Liu, Myeonghoon Han, Emily Stoneburner, Stephanie Makdissi, Francesca Di Cara, Aaron Ring, and Norbert Perrimon. Phage-displayed synthetic library and screening platform for nanobody discovery. eLife, 14:RP105887, 2025. doi: 10.7554/eLife.105887.

[33] Weijie Zhang, Hao Wang, Nan Feng, Yifeng Li, Jijie Gu, and Zhuozhi Wang. Developability assessment at early-stage discovery to enable development of antibody-derived therapeutics. Antibody Therapeutics, 6(1):13–29, 2023. doi: 10.1093/abt/tbac029.

[34] Matthew N. Mendoza, Mike Jian, Moeko T. King, and Cory L. Brooks. Role of a noncanonical disulphide bond in the stability, affinity, and flexibility of a vhh specific for the listeria virulence factor inlb. Protein Science, 29(4):1004–1017, 2020. doi: 10.1002/pro.3831.

[35] Monica L. Fernández-Quintero et al. On the humanization of VHHs: Prospective case studies, experimental and computational characterization of structural determinants for functionality. Protein Science, 2024. doi: 10.1002/pro.5176.

[36] Aubin Ramon, Niccolò Frassetto Haowen Zhao, Xing Xu, Matthew Greenig, Shimobi Onuoha, and Pietro Sormanni. Deep learning assessment of nativeness and pairing likelihood for antibody and nanobody design with abnativ2. bioRxiv, page 2025.10.31.685806, 2025. doi: 10.1101/2025.10.31.685806.

[37] James Dunbar, Konrad Krawczyk, Jinwoo Leem, Terry Baker, Angelika Fuchs, Guy Georges, Jiye Shi, and Charlotte M. Deane. Sabdab: the structural antibody database. Nucleic Acids Research, 42(D1):D1140–D1146, 2014. doi: 10.1093/nar/gkt1043.

[38] Sen Zheng. Navigating the unstructured by evaluating alphafold’s efficacy in predicting missing residues and structural disorder in proteins. PLoS ONE, 20(3):e0313812, 2025. doi: 10.1371/journal.pone.0313812.

[39] Nathaniel R. Bennett, Joseph L. Watson, Robert J. Ragotte, Andrew J. Borst, DéJenaé L. See, Connor Weidle, Riti Biswas, Yutong Yu, Ellen L. Shrock, Russell Ault, Philip J. Y. Leung, Buwei Huang, Inna Goreshnik, John Tam, Kenneth D. Carr, Benedikt Singer, Cameron Criswell, Basile I. M. Wicky, Dionne Vafeados, Mariana Garcia Sanchez, Ho Min Kim, Susana Vázquez Torres, Sidney Chan, Shirley M. Sun, Timothy T. Spear, Yi Sun, Keelan O’Reilly, John M. Maris, Nikolaos G. Sgourakis, Roman A. Melnyk, Chang C. Liu, and David Baker. Atomically accurate de novo design of antibodies with rfdiffusion. Nature, 649:183–193, 2025. doi: 10.1038/s41586-025-09721-5.

[40] Thomas H. Olsen, Frederick Boyles, and Charlotte M. Deane. Observed antibody space: A diverse database of cleaned, annotated, and translated unpaired and paired antibody sequences. Protein Science, 31(1):141–146, 2022. doi: 10.1002/pro.4205.

[41] Pawel Dudzic, Dawid Chomicz, Jarosław Konćzak, Tadeusz Satława, Bartosz Janusz, Sonia Wrobel, Tomasz Gawłowski, Igor Jaszczyszyn, Weronika Bielska, Samuel Demharter, Roberto Spreafico, Lukas Schulte, Kyle Martin, Stephen R. Comeau, and Konrad Krawczyk. Large-scale data mining of four billion human antibody variable regions reveals convergence between therapeutic and natural antibodies that constrains search space for biologics drug discovery. mAbs, 16(1):2361928, 2024. doi: 10.1080/19420862.2024.2361928.

[42] Martin Steinegger and Johannes Söding. Mmseqs2 enables sensitive protein sequence searching for the analysis of massive data sets. Nature Biotechnology, 35(11):1026–1028, 2017. doi: 10.1038/nbt.3988. URL https://www.nature.com/articles/nbt.3988.

[43] Monica L. Fernández-Quintero, Janik Kokot, Franz Waibl, Anna-Lena M. Fischer, Patrick K. Quoika, Charlotte M. Deane, and Klaus R. Liedl. Challenges in antibody structure prediction. mAbs, 15(1):2175319, 2023. doi: 10.1080/19420862.2023.2175319. URL https://pubmed.ncbi.nlm.nih.gov/36775843/.

[44] Montader Ali, Mateusz Jaskolowski, Matthew Greenig, Mia Crnogaj, Eva Smorodina, Haowen Zhao, Victor Greiff, and Pietro Sormanni. Improving nanobody structure prediction with self-distillation. bioRxiv, 2025. doi: 10.64898/2025.12.01.691162.

[45] Brennan Abanades, Wing Ki Wong, Fergus Boyles, Guy Georges, Alexander Bujotzek, and Charlotte M. Deane. Immunebuilder: Deep-learning models for predicting the structures of immune proteins. Communications Biology, 2023. doi: 10.1038/s42003-023-04927-7.

[46] Margarida C. Simões, Joana S. Cristóvão, Els Pardon, Jan Steyaert, Günter Fritz, and Cláudio M. Gomes. Functional modulation of rage activation by multimeric s100b using single-domain antibodies. Journal of Biological Chemistry, 300(12):107983, 2024. doi: 10.1016/j.jbc.2024.107983.

[47] Josh Abramson et al. Accurate structure prediction of biomolecular interactions with AlphaFold 3. Nature, 2024. doi: 10.1038/s41586-024-07487-w.

[48] Frédéric A. Dreyer, Jan Ludwiczak, Karolis Martinkus, Brennan Abanades, Robert G. Alberstein, Pan Kessel, Pranav Rao, Jae Hyeon Lee, Richard Bonneau, Andrew M. Watkins, and Franziska Seeger. Conformation-aware structure prediction of antigen-recognizing immune proteins. mAbs, 18(1):2602217, 2025. doi: 10.1080/19420862.2025.2602217. URL https://www.tandfonline.com/doi/full/10.1080/19420862.2025.2602217.

[49] Xinhao Wang, Lu Zhang, Yao Zhang, Jiaguo Li, Wenfeng Xu, and Weimin Zhu. Distinct types of vhhs in alpaca. Frontiers in Immunology, 15:1447212, 2024. doi: 10.3389/fimmu.2024.1447212. URL https://www.ncbi.nlm.nih.gov/pmc/articles/PMC11588638/.

[50] Robert J. Hoey, Hyeyoung Eom, and James R. Horn. Structure and development of single domain antibodies as modules for therapeutics and diagnostics. Experimental Biology and Medicine, 244(17):1568–1576, 2019. doi: 10.1177/1535370219881129. URL https://www.ncbi.nlm.nih.gov/pmc/articles/PMC6920669/.

[51] Erik Swanson, Michael Nichols, Supriya Ravichandran, and Pierce Ogden. mber: Controllable de novo antibody design with million-scale experimental screening. bioRxiv, 2025. doi: 10.1101/2025.09.26.678877. URL https://www.biorxiv.org/content/10.1101/2025.09.26.678877.

[52] Aubin Ramon, Montader Ali, Misha Atkinson, Alessio Saturnino, Kieran Didi, Cristina Visentin, Stefano Ricagno, Xing Xu, Matthew Greenig, and Pietro Sormanni. Assessing antibody and nanobody nativeness for hit selection and humanization with abnativ. Nature Machine Intelligence, 6:74–91, 2024. doi: 10.1038/s42256-023-00778-3. URL https://www.nature.com/articles/s42256-023-00778-3.

[53] Aubin Ramon, Mingyang Ni, Olga Predeina, Rebecca Gaffey, and Pietro Sormanni. Prediction of protein biophysical traits from limited data: a case study on nanobody thermostability through nanomelt. mAbs, 17(1):2442750, 2025. doi: 10.1080/19420862.2024.2442750.

[54] E. Gordon and C. M. Deane. The therapeutic nanobody profiler: characterising and predicting nanobody developability to improve therapeutic design. bioRxiv, 2025. doi: 10.1101/2025.08.11.669635. URL https://www.biorxiv.org/content/10.1101/2025.08.11.669635.

[55] Luis S. Mille-Fragoso, John N. Wang, Claudia L. Driscoll, Haoyu Dai, Talal M. Widatalla, Xiaowei Zhang, Brian L. Hie, and Xiaojing J. Gao. Efficient generation of epitope-targeted de novo antibodies with germinal. bioRxiv, 2025. doi: 10.1101/2025.09.19.677421. URL https://www.biorxiv.org/content/10.1101/2025.09.19.677421v2.

[56] Mahmoud Ganji, Pooria Safarzadeh Kozani, and Fatemeh Rahbarizadeh. Characterization of novel cd19-specific vhhs isolated from a camelid immune library by phage display. Journal of Translational Medicine, 21(1):891, 2023. doi: 10.1186/s12967-023-04524-6. URL https://doi.org/10.1186/s12967-023-04524-6.

[57] Vishakha Singh, Mandar Bhutkar, Shweta Choudhary, Sanketkumar Nehul, Rajesh Kumar, Jitin Singla, Pravindra Kumar, and Shailly Tomar. Structure-guided mutations in cdrs for enhancing the affinity of neutralizing sars-cov-2 nanobody. Biochemical and Biophysical Research Communications, 734:150746, 2024. doi: 10.1016/j.bbrc.2024.150746. URL https://pubmed.ncbi.nlm.nih.gov/39366179/.

[58] Patrick Kunz, Katinka Zinner, Norbert Mücke, Tanja Bartoschik, Serge Muyldermans, and Jörg D. Hoheisel. Influence of the conserved disulphide bond on the folding and stability of camelid vhh domains. Journal of Molecular Biology, 430(3):277–295, 2018. doi: 10.1016/j.jmb.2017.11.007.

[59] Jochen Govaert, Mireille Pellis, Nick Deschacht, Cécile Vincke, Katja Conrath, Serge Muyldermans, and Dirk Saerens. Dual beneficial effect of interloop disulphide bond for single domain antibody fragments. Journal of Biological Chemistry, 287(3):1970–1979, 2012. doi: 10.1074/jbc.M111.242818. URL https://pubmed.ncbi.nlm.nih.gov/22128183/.

[60] Monica L. Fernández-Quintero, Johannes R. Loeffler, Lisa M. Bacher, Franz Waibl, Clarissa A. Seidler, and Klaus R. Liedl. Local and global rigidification upon antibody affinity maturation. Frontiers in Molecular Biosciences, 7:182, 2020. doi: 10.3389/fmolb.2020.00182. URL https://www.ncbi.nlm.nih.gov/pmc/articles/PMC7426445/.

[61] Yuning Shen, Lihao Wang, Huizhuo Yuan, Yan Wang, Bangji Yang, and Quanquan Gu. Confrover: Simultaneous modeling of protein conformation and dynamics via autoregression. arXiv, 2025. doi: 10.48550/arXiv.2505.17478. URL https://arxiv.org/abs/2505.17478v2.

[62] Sarah Lewis, Tim Hempel, José Jiménez-Luna, Michael Gastegger, Yu Xie, Andrew Y. K. Foong, Victor García Satorras, Osama Abdin, Bastiaan S. Veeling, Iryna Zaporozhets, Yaoyi Chen, Soojung Yang, Adam E. Foster, Arne Schneuing, Jigyasa Nigam, Federico Barbero, Vincent Stimper, Andrew Campbell, Jason Yim, Marten Lienen, Yu Shi, Shuxin Zheng, Hannes Schulz, Usman Munir, Roberto Sordillo, Ryota Tomioka, Cecilia Clementi, and Frank Noé. Scalable emulation of protein equilibrium ensembles with generative deep learning. Science, 389(6761):eadv9817, 2025. doi: 10.1126/science.adv9817. URL https://www.science.org/doi/10.1126/science.adv9817.

[63] James P. Roney, Chenxi Ou, and Sergey Ovchinnikov. Protein diffusion models as statistical potentials. bioRxiv, 2025. doi: 10.64898/2025.12.09.693073. URL https://www.biorxiv.org/content/10.64898/2025.12.09.693073v1.

[64] A. Greenshields-Watson, O. Vavourakis, F. C. Spoendlin, M. Cagiada, and C. M. Deane. Challenges and compromises: Predicting unbound antibody structures with deep learning. Current Opinion in Structural Biology, 90:102983, 2025. ISSN 0959-440X. doi: 10.1016/j.sbi.2025.102983. URL https://www.sciencedirect.com/science/article/pii/S0959440X25000016.

[65] Monica L. Fernández-Quintero, Johannes R. Loeffler, Johannes Kraml, Ursula Kähler, Anna S. Kamenik, and Klaus R. Liedl. Characterizing the diversity of the cdrh3 loop conformational ensembles in relationship to antibody binding properties. Frontiers in Immunology, 9:3065, 2019. doi: 10.3389/fimmu.2018.03065. URL https://www.frontiersin.org/articles/10.3389/fimmu.2018.03065/full.

[66] Wei Wang, Wei Ye, Qing Yu, Cheng Jiang, Jian Zhang, and Rui Luo. Conformational selection and induced fit in specific antibody and antigen recognition: Spe7 as a case study. The Journal of Physical Chemistry B, 117(17):4912–4923, 2013. doi: 10.1021/jp4010967. URL https://pubs.acs.org/doi/10.1021/jp4010967.

[67] Chu’nan Liu, Lilian M Denzler, Oliver E.C. Hood, and Andrew C.R. Martin. Do antibody cdr loops change conformation upon binding? mAbs, 16(1):2322533, 2024. doi: 10.1080/19420862.2024.2322533. URL https://www.ncbi.nlm.nih.gov/pmc/articles/PMC10939163/.

[68] Annemarie Honegger and Andreas Plückthun. Yet another numbering scheme for immunoglobulin variable domains: An automatic modeling and analysis tool. Journal of Molecular Biology, 309(3):657–670, 2001. ISSN 0022-2836. doi: 10.1006/jmbi.2001.4662.

[69] Simon Mitternacht. Freesasa: An open source c library for solvent accessible surface area calculations. F1000Research, 5:189, 2016. doi: 10.12688/f1000research.7931.1. URL https://f1000research.com/articles/5-189/v1.

[70] Matthew Z. Tien, Austin G. Meyer, Dariya K. Sydykova, Stephanie J. Spielman, and Claus O. Wilke. Maximum allowed solvent accessibilites of residues in proteins. PLoS ONE, 8(11):e80635, 2013. doi: 10.1371/journal.pone.0080635.

[71] F. Pedregosa, G. Varoquaux, A. Gramfort, V. Michel, B. Thirion, O. Grisel, M. Blondel, P. Prettenhofer, R. Weiss, V. Dubourg, J. Vanderplas, A. Passos, D. Cournapeau, M. Brucher, M. Perrot, and E. Duchesnay. Scikit-learn: Machine learning in Python. Journal of Machine Learning Research, 12:2825–2830, 2011.

[72] Henry Kenlay, Frédéric A. Dreyer, Daniel Cutting, Daniel Nissley, and Charlotte M. Deane. Abodybuilder3: Improved and scalable antibody structure predictions. Bioinformatics, 40(10):btae576, 2024. doi: 10.1093/bioinformatics/btae576. URL https://academic.oup.com/bioinformatics/advance-article/doi/10.1093/bioinformatics/btae576/7810444.

[73] Gustaf Ahdritz, Nazim Bouatta, Christina Floristean, Sachin Kadyan, Daniel Berenberg O’Donnell, Ian Fisk, Niccolò Zanichelli Bo Zhang, Arkadiusz Nowaczynski, Bei Wang, Marta M. Stepniewska-Dziubinska, Shang Zhang, Adegoke A. Ojewole, Murat Efe Guney, Stella Biderman, Andrew M. Watkins, Stephen Ra, Pablo Ribalta Lorenzo, Lucas Nivon, Brian D. Weitzner, Yih-En Andrew Ban, Shiyang Chen, Minjia Zhang, Conglong Li, Shuaiwen Leon Song, Yuxiong He, Peter K. Sorger, Emad Mostaque, Zhao Zhang, Richard Bonneau, Mohammed AlQuraishi, et al. Openfold: retraining alphafold2 yields new insights into its learning mechanisms and capacity for generalization. Nature Methods, 21(8):1514–1524, 2024. doi: 10.1038/s41592-024-02272-z.

[74] James A. Maier, Carmenza Martinez, Koushik Kasavajhala, Lauren Wickstrom, Kevin E. Hauser, and Carlos Simmerling. ff14sb: Improving the accuracy of protein side chain and backbone parameters from ff99sb. Journal of Chemical Theory and Computation, 11(8):3696–3713, 2015. doi: 10.1021/acs.jctc.5b00255. URL https://doi.org/10.1021/acs.jctc.5b00255.

[75] Peter Eastman, Raimondas Galvelis, R. P. Peláez, Charlles R. A. Abreu, Stephen E. Farr, Emilio Gallicchio, Anton Gorenko, Michael M. Henry, Frank Hu, Jing Huang, Andreas Krämer, Julien Michel, Joshua A. Mitchell, Vijay S. Pande, João P. G. L. M. Rodrigues, Ivanka Savic, Andrew C. Simmonett, Sukrit Singh, Jason Swails, Philip Turner, Yuanqing Wang, Ivy Zhang, John D. Chodera, Gianni de Fabritiis, and Thomas Edward Markland. Openmm 8: Molecular dynamics simulation with machine learning potentials. Journal of Physical Chemistry B, 128 (1):109–116, 2023. doi: 10.1021/acs.jpcb.3c06662. URL https://pubs.acs.org/doi/10.1021/acs.jpcb.3c06662. Published online December 28 2023.

[76] Ming-an Sun, Yejun Wang, Qing Zhang, Yiji Xia, Wei Ge, and Dianjing Guo. Prediction of reversible disulphide based on features from local structural signatures. BMC Genomics, 18:279, 2017. doi: 10.1186/s12864-017-3668-8. URL https://www.ncbi.nlm.nih.gov/pmc/articles/PMC5379614/.

[77] Milot Mirdita, Konstantin Schütze, Yoshitaka Moriwaki, Lim Heo, Sergey Ovchinnikov, and Martin Steinegger. Colabfold: making protein folding accessible to all. Nature Methods, 19 (6):679–682, 2022. doi: 10.1038/s41592-022-01488-1. URL https://www.nature.com/articles/s41592-022-01488-1.

[78] Richard Evans, Michael O’Neill, Alexander Pritzel, Natasha Antropova, Andrew W. Senior, Tim Green, Augustin Žídek, Robert Bates, Sam Blackwell, Jason Yim, Olaf Ronneberger, Sebastian Bodenstein, Michal Zielinski, Alex Bridgland, Andrey Potapenko, Andrew Cowie, Kathryn Tunyasuvunakool, Rishub Jain, Ellen Clancy, Pushmeet Kohli, John Jumper, and Demis Hassabis. Protein complex prediction with alphafold-multimer. bioRxiv, 2021. doi: 10.1101/2021.10.04.463034. URL https://www.biorxiv.org/content/10.1101/2021.10.04.463034v2.

[79] S Basu and B Wallner. Dockq: A quality measure for protein-protein docking models. PLOS ONE, 11(8):e0161879, 2016. doi: 10.1371/journal.pone.0161879. URL https://journals.plos.org/plosone/article?id=10.1371/journal.pone.0161879.

